# DiffSegR: An RNA-Seq data driven method for differential expression analysis using changepoint detection

**DOI:** 10.1101/2023.06.05.543691

**Authors:** Arnaud Liehrmann, Etienne Delannoy, Alexandra Launay-Avon, Elodie Gilbault, Olivier Loudet, Benoît Castandet, Guillem Rigaill

## Abstract

To fully understand gene regulation, it is necessary to have a thorough understanding of both the transcriptome and the enzymatic and RNA-binding activities that shape it. While many RNA-Seq-based tools have been developed to analyze the transcriptome, most only consider the abundance of sequencing reads along annotated patterns (such as genes). These annotations are typically incomplete, leading to errors in the differential expression analysis. To address this issue, we present DiffSegR - an R package that enables the discovery of transcriptome-wide expression differences between two biological conditions using RNA-Seq data. DiffSegR does not require prior annotation and uses a multiple changepoints detection algorithm to identify the boundaries of differentially expressed regions in the per-base log2 fold change. In a few minutes of computation, DiffSegR could rightfully predict the role of chloroplast ribonuclease Mini-III in rRNA maturation and chloroplast ribonuclease PNPase in (3’/5’)-degradation of rRNA, mRNA, and tRNA precursors as well as intron accumulation. We believe DiffSegR will benefit biologists working on transcriptomics as it allows access to information from a layer of the transcriptome overlooked by the classical differential expression analysis pipelines widely used today. DiffSegR is available at https://aliehrmann.github.io/DiffSegR/index.html.

## INTRODUCTION

It has long been recognized that transcriptomes largely surpass genomes in complexity (1). Besides alternative use of transcription initiation sites, most of the transcript diversity can be ascribed to post-transcriptional modifications, including RNA splicing, processing, alternative polyadenylation, editing or base modification (2). Although the advent of the transcriptomics revolution has allowed an unprecedented understanding of this transcript diversity, the combinatorial nature and very large number of variations is still an analytical challenge (3, 4). Moreover, because most strategies for RNA-Seq analysis rely on incomplete transcriptomic variant annotations, meaningful variations may currently be overlooked (5). This is a major issue for biological interpretation as illustrated by the crucial role played in disease development by poorly annotated non coding elements like 5’ and 3’ UTRs (6–9).

As a consequence, there is a massive effort to improve transcriptomic annotations with the help of the third generation (long-read) sequencing technologies from Oxford Nanopore Technologies or Pacific Bioscience. Long RNA-Seq reads may cover an entire RNA isoform from start to end, directly illustrating the exon structure, splicing patterns and UTR composition (10–12). They carry the promise to go beyond the limits of full-length transcript assembly, which is notoriously prone to error (13, 14). Although such a strategy can double the number of referenced transcripts for a model organism (15), reaching an exhaustive description of a transcriptome is arguably a Sisyphean task (5, 16, 17).

Because most RNA-Seq experiments aim at identifying RNA processes that vary between two biological conditions (WT vs mutant or control vs stress, for example), an alternative to this issue is to identify portions of the transcriptome that vary between both experimental conditions (differentially expressed regions or DERs) directly from the RNA-Seq data, without relying on annotations and bypassing assembly altogether. This is performed by a class of methods sometimes referred to as identify-then-annotate tools (18). Their gold standard is to be both highly specific (i.e. able to merge adjacent non-DERs) and highly sensitive (i.e. able to discriminate between adjacent DERs, in particular between up and down DERs). To do so, various methods summarized in Figure 1 (19–22) address a well-defined statistical problem known as multiple changepoints detection, or segmentation problem. This has been a long-standing problem in the field of genomic series analysis (23–27). To identify DERs, current identify-then-annotate tools mainly vary in the signal they segment and in the way they segment it (Figure 1).

**Figure 1:**
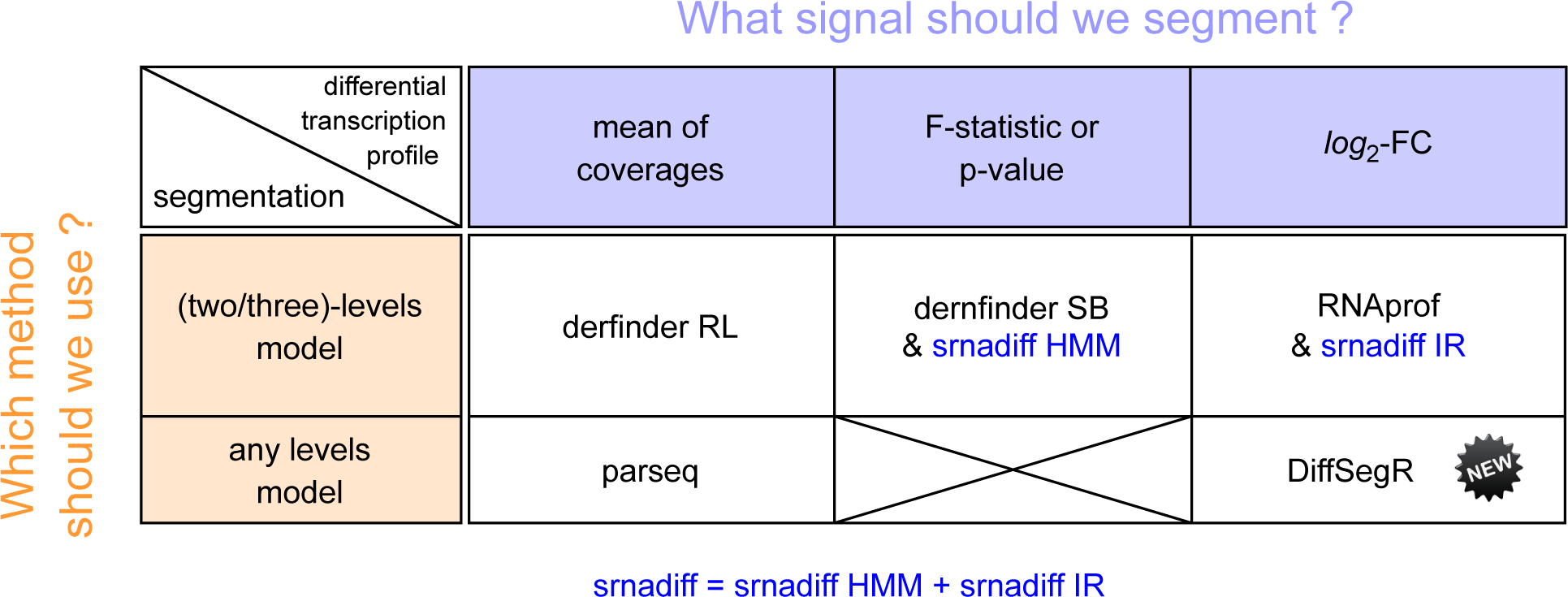
State-of-the-art of Identify-then-annotate methods for detecting differentially expressed regions (DERs) in RNA-Seq data. The methods included in this figure - srnadiff, srnadiff IR, srnadiff HMM (19), derfinder SB, derfinder RL(22), RNAprof (21), parseq (20), and DiffSegR - belong to a class of methods known as identified-then-annotate, which enable the identification of DERs directly from RNA-Seq data without relying on annotations or assembly. To identify DERs, these methods address a well-defined statistical problem known as multiple changepoints detection or segmentation problem. The methods vary in the signal they segment and the way they segment it. For example, srnadiff merges the results of a three-level segmentation model on the per-base log2 fold-change (srnadiff IR) and a two-level segmentation model on the per- base p-value (srnadiff HMM). Similarly, derfinder SB, and derfinder RL implement a two- level segmentation model on the per-base p-value, per-base F-statistic (similar to per-base p- value), and the mean of coverages, respectively. RNAprof implements a three level segmentation model on the per-base log2 fold-change. parseq segments the mean of coverages without assuming the number of levels. Finally, DiffSegR introduces a new strategy to identify DERs by segmenting the per-base log2 fold-change without assuming the number of levels. All the methods except parseq assessed the found DERs using DESeq2 (29).

Here, we introduced DiffSegR, an R package that uses a new strategy for delineating the boundaries of DERs. It segments the per-base log2 fold change (log2-FC) using FPOP, a method designed to identify changepoints in the mean of a Gaussian signal (28). Intuitively, the per-base log2-FC is a measure that scales with the intensity of the transcription differences at each genomic position between the two compared biological conditions. Expression differences are then statistically assessed for each region using the negative binomial generalized linear model of DESeq2 (29) and the outputs can be visualized in IGV (30).

DiffSegR and competitor methods (Figure 1) were compared on two plant RNA-Seq datasets that were previously used in combination with molecular biology techniques to decipher the roles of the chloroplast ribonucleases PNPase and Mini-III (31, 32). DiffSegR was the only method able to retrieve all the segments known to differentially accumulate outside of the annotated genic regions (i.e. 3’ and 5’ extensions, anti-sense accumulation). Moreover, it is the only method predicting the overaccumulation of intronic regions on the plastome-scale in the PNPase mutant. Globally, DiffSegR better captures multiple trends of differences within DERs while being more parsimonious in non-DERs than its competitors.

We anticipate DiffSegR will be an important tool in providing an in-depth description of local or regional transcript variations within RNA-seq libraries from two biological conditions, especially when studying RNA processes located outside of the annotated coding sequences, like RNA processing, trimming or splicing.

## MATERIALS AND METHODS

### DiffSegR segmentation model

#### Differential transcription profile

DiffSegR builds the coverage profiles indexed on *n* genomic positions from the BAM files provided by the user. The coverage profile for replicate *r* of biological condition *j* is noted 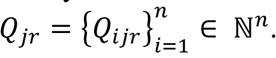. By default we propose to compute *Q*_ijr_ using the geometric mean of the number of 5’ and 3’ end of the reads overlapping position *i*, denoted *Qijr5’* and *Qijr3’* . Formally:

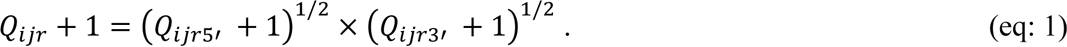

We describe alternative approaches that use either the full length or the 5’ or 3’ end of the reads, and compare them with our geometric mean heuristic in Notes S1-3. DiffSegR then builds the differential transcription profile between the biological conditions (named *1* and *2* hereafter) using a log_2_-FC per-base transformation because it scales with the intensity of the transcriptional differences between conditions *1* and *2*. The log2-FC at the *i* ^th^ genomic position (denoted *Yi*) is given by

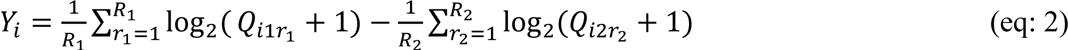

where *R1* and *R^2^* stand for the number of replicates in condition *1* and *2*, respectively.

#### Segmentation model

We consider *D* changepoints *τ1<…< τD* within the range 1 and *n* - 1. These changepoints correspond to unknown positions along the genome where a shift in the mean of the per-base log2-FC (eq: 2) is observed. We adopt the convention that *τ0* = 0 and *τ*|*τ*| = *n*. These changepoints define |*τ*| = *D* + 1 distinct segments. The *j^th^*segment includes the data ⟧*τj- 1,τj*⟧={*τj-1+*1,…,*τj*}. Each segment is premised on the assumption that the *Yi* therein are independent and follow the same Gaussian distribution, with a mean *µj* specific to that segment and a common variance *σ^2^*. Expressed mathematically, we have:

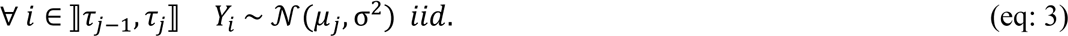

#### Estimation of the segment

The parameters of the model (eq: 3), including *τ1<…< τD*, can be estimated using penalized maximum likelihood inference. To achieve this, DiffSegR uses the FPOP algorithm (28) (a dynamic programming algorithm that implements functional pruning techniques) which solves the inference problem exactly (see below). For many profiles lengths the computation time of FPOP is log-linear allowing for the segmentation of large data (*10^6^<n<10^7^*) in a matter of seconds. The number of changepoints estimated by FPOP is a decreasing function of the penalty *λσ^2^* log(*n*). The constant *λ* is a hyperparameter that can be adjusted by the user. A good starting point, based on theoretical arguments (33) and simulations (34), is to set *λ* = 2. The variance *σ^2^* is estimated on the data using the unbiased sample variance estimator.

#### FPOP

Informally, the idea of the FPOP algorithm is to consider the penalized maximum likelihood of the data from observation 1 to *t* as a function of the parameter of the last segment. This idea is referred to as ’functional pruning.’ In the Gaussian case, the resulting function is piecewise quadratic. For a new observation at time *t*+1, it is possible to efficiently update this function (that is, compute the penalized maximum likelihood function from observation 1 to *t*+1) using a formula similar to that of the Viterbi algorithm. This formula is applied piece by piece, that is by intervals. At each step, the algorithm searches for the best possible value of the parameter of the last segment to maximize the penalized likelihood.

### Normalization

To account for differences in the total number of sequenced reads per sample, we assume that the mean of the coverage *µijr* is composed of a sample-specific size factor *sjr* and a parameter *qijr* proportional to the expected true concentration of transcripts overlapping position *i* in replicate *r* of condition *j* verifying *μijr = sjr qijr* (29, 35, 36). As the coverage (eq: 1), the per-base log2-FC (eq: 2) depends on sample-specific size factors. One can show that the non-normalized and normalized per-base log2-FC are linked together by an offset denoted *ρ* such that

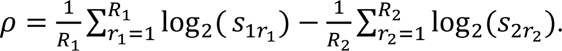

For a given penalty the output of FPOP is shift invariant. That is if the data is shifted by a given value the returned changepoints will be the same. Therefore the segmentation returned by DiffSegR does not depend on the knowledge of the normalization factors. This is a key difference with threshold based methods (e.g. srnadiff IR, srnadiff HMM, RNAprof, derfinder RL, derfinder SB).

We acknowledge that when taking into account the offset to the logarithms (+1) in the per-base log2-FC, the previous argument is approximately true for large counts but does not hold for small counts.

### Overview of the DiffSegR package

DiffSegR is implemented as an R package (www.R-project.org/) and can be found on GitHub (https://github.com/aLiehrmann/DiffSegR) with the installation procedure as well as a vignette with functional examples. The package implements the four steps of a conventional pipeline for identify-then-annotate methods (Figure 2).

**Figure 2:**
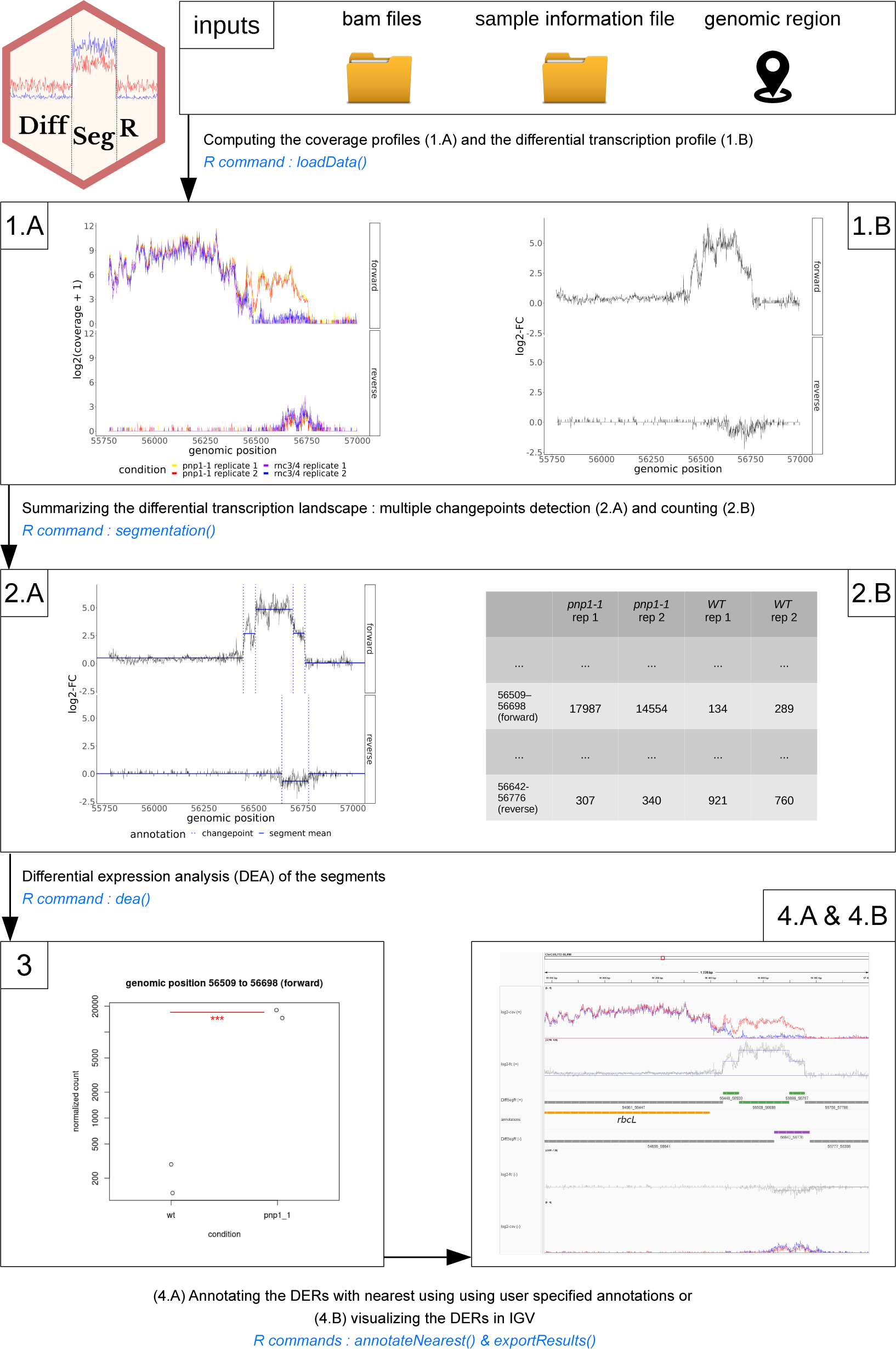
Schematic representation of the DiffSegR pipeline. The DiffSegR pipeline consists of four major steps: **(1)** Computing the coverage profiles and the differential transcription profile from BAMs. The *loadData* function creates coverage profiles from user-specified BAM files and a genomic region. (1.A) It produces one profile per strand for each replicate of both biological conditions. (1.B) The function then calculates the per-base log2 fold-change (log2-FC) based on the coverage profiles. **(2)** Summarizing the differential transcription landscape. (2.A) The *segmentation* function applies FPOP to the per-base log2-FC of each strand to identify segment boundaries, known as changepoints. (2.B) Then the *featurecounts* program is used to assign mapped reads to segments, resulting in a count matrix. **(3)** Differential expression analysis (DEA). The *dea* function uses DESeq2 to test the difference in average expression between the two compared biological conditions for each segment. **(4)** Annotating and visualizing the differentially expressed regions (DERs). (4.A) The *annotateNearest* function annotates DERs using user-specified gff3 or gtf format annotations. In parallel, (4.B) the *exportResults* function saves DERs, not-DERs, segmentation, the mean of coverage profiles from both biological conditions, and per-base log2-FC information in formats compatible with genome viewers like IGV. An IGV session in XML format allows loading all tracks with one click, providing a user-friendly way to visualize and interpret DiffSegR results.

#### Step 1: Computing the coverage profiles and the differential transcription profile from BAMs

The *loadData* function builds coverage profiles from BAM files within a locus specified by the user. If the reads are stranded, the function builds one coverage profile per strand for each replicate of both compared biological conditions. By default the heuristic used to compute coverage profiles is the geometric mean of the 5’ and 3’ profiles (eq: 1). Alternative approaches use either the full length or the 5’ or 3’ end of the reads (Notes S1). *loadData* then converts the coverage profiles into the per-base log2-FC (eq: 2) (one per strand) using the reference biological condition specified by the user as the denominator. The function returns the coverage profiles and the differential transcription profile as a list of run-length encoded objects.

#### Step 2: Summarizing the differential transcription landscape

The *segmentation* function uses FPOP (28) on the per-base log2-FC of both strands to identify the segment’s boundaries (or changepoints). The number of returned segments is controlled by the hyperparameter *λ* specified by the user. The segments are stored as GenomicRanges object and the *segmentation* function finally uses *featurecounts* (44) to assign them the mapped reads from each replicate of each biological condition. By default a read is allowed to be assigned to every segment it overlaps with. The segments and the associated count matrix are returned as a SummarizedExperiment object.

#### Step 3: Differential expression analysis (DEA)

The *dea* function uses DESeq2 (29) to test the difference in average expression between the two compared biological conditions for every segment. The resulting p-values are then adjusted using a Benjamini-Hochberg (BH) procedure to control the false discovery rate (FDR), which is a common approach in DEA. However, this approach does not guarantee that the proportion of false discoveries (FDP) will be upper bounded, and there is no statistical guarantee on the number of false discoveries in subsets of segments selected using FDR thresholding. For example, while a widespread practice in DEA is to select a subset of segments whose absolute log2-FC passes a threshold it can potentially result in an inflated FDR. To address these limitations, the *dea* function can also call a post-hoc inference procedure that provides guarantees on the FDP in arbitrary segment selections (42). Finally, *dea* returns the user-provided SummarizedExperiment object augmented with the outcome of the DEA.

#### Step 4.A: Annotating the Differentially expressed regions (DERs)

The *annotateNearest* function annotates the DERs found during the DEA using user specified annotations in the gff3 or gtf format. Seven classes of labels translate the relative positions of the DER and its closest annotation(s): antisense, upstream, downstream, inside, overlapping 3’, overlapping 5’ and overlapping both 5’ and 3’. These labels allow users to easily understand the relationships between the DERs and their nearest annotations, and to analyze the potential functional significance of the DERs in the context of the annotated genomic features.

#### Step 4.B: Visualizing the DERs

The *exportResults* function saves the DERs, not-DERs, segmentation, the mean of coverage profiles from both biological conditions and per-base log2-FC information, for both strands, in formats readable (bed, gff3) by genome viewers like the Integrative Genome Viewer (IGV) (30). For IGV, *exportResults* also creates a session in xml format that allows loading all tracks in one click. This provides a convenient way to save and visualize the results of the differential expression analysis, allowing a user-friendly exploration and interpretation of the data generated by the DEA. An example of the graphical output obtained with DiffSegR is displayed in Figure 3.

**Figure 3:**
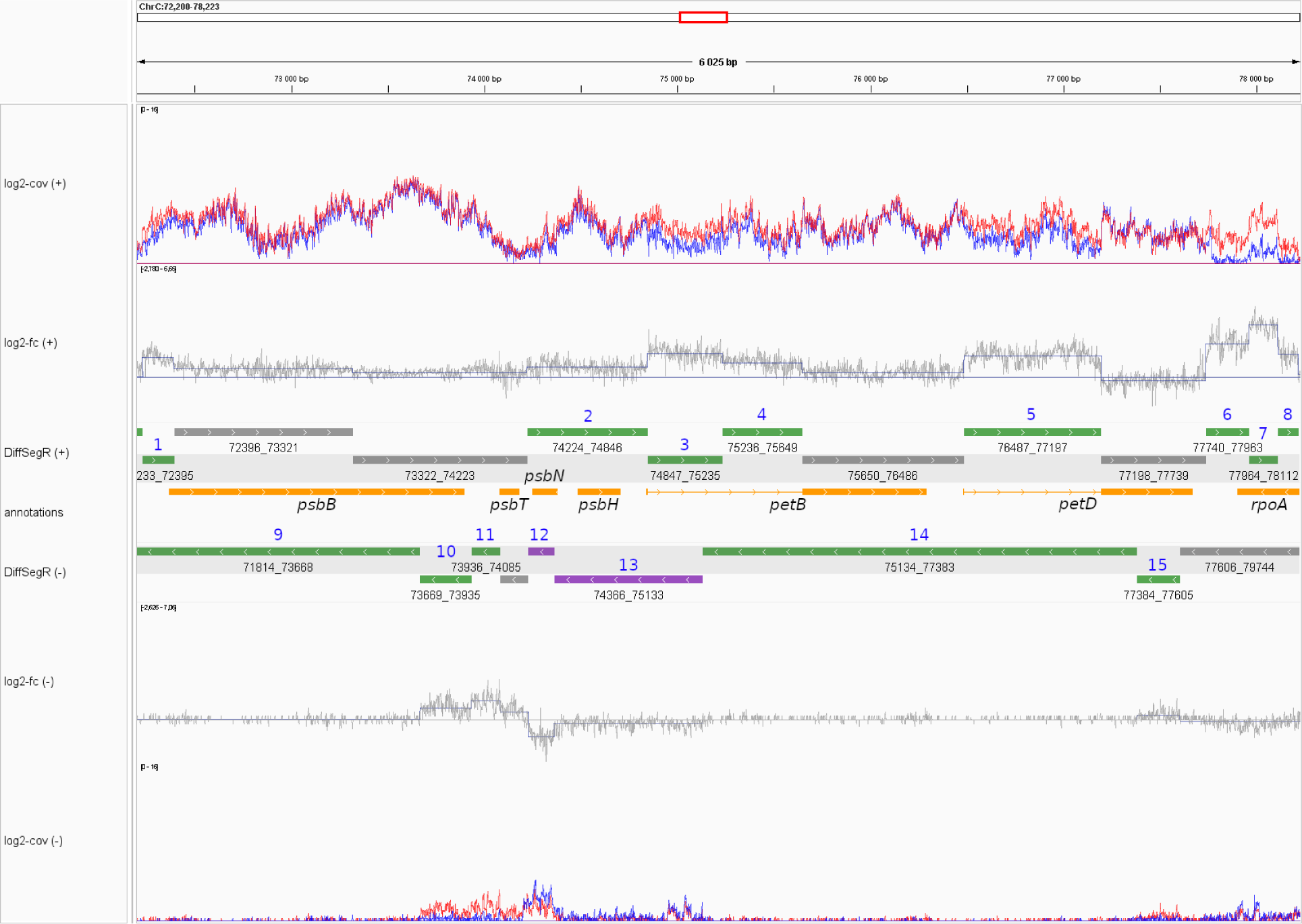
DiffSegR analysis of the *psbB-psbT-psbN-psbH-petB-petD* gene cluster in the *pnp1-1* dataset. The tracks from top to bottom represent: (log2-Cov (+)) the mean of coverages on the log2 scale for the forward strand in both biological conditions of interest, with the blue line representing the *WT* condition and the red line representing the *pnp1-1* condition; (log2-FC (+)) the per-base log2-FC between *pnp1-1* (numerator) and WT (denominator) for the forward strand. The straight horizontal line represents the zero indicator. When the per-base log2-FC is above or below the zero indicator line, it suggests up-regulation or down- regulation, respectively, in *pnp1-1* compared to *WT.* The changepoint positions are indicated by vertical blue lines, and the mean of each segment is shown by horizontal blue lines connecting two changepoints; (DiffSegR (+)) the differential expression analysis results for segments identified by DiffSegR on the forward strand are presented as follows: up-regulated regions are depicted in green, down-regulated regions in purple, and non-differentially expressed regions in gray; (annotations) the genes provided by users for interpretations. Symmetrically, the remaining tracks correspond to the same data on the reverse strand. DiffSegR finds 8 up-regulated DERs on the forward strand (IDs 1 to 8), 5 up-regulated DERs on the reverse strand (IDs 9 to 11, 14 and 15), and 2 down-regulated DERs on the reverse strand (IDs 12 and 13). Table 1 provides a summary of the molecular validations published for the DERs identified in the *psbB* gene cluster through DiffSegR analysis. The bedGraph and gff3 files used to generate the tracks and the xml file used to load them in IGV were created using the *exportResults* function of the DiffSegR R package. The session was loaded in IGV 2.12.3 for Linux.

**Table 1:** DERs identified by DiffSegR within the gene cluster *psbB-psbT-psbN-psbH-petB- petD* in *pnp1-1* dataset. Most DERs are supported by molecular validations described in the literature. Up is for up-regulated and down for down-regulated.

| strand | positions | DiffSegR<br>result | genomic context | ID | validation |
| --- | --- | --- | --- | --- | --- |
| forward | 72,233-72,395 | up | *psbB* 5' ends | 1 | (45) |
| forward | 74,224-74,846 | up | *psbH*; antisense to *psbN* | 2 | (37, 46–48) |
| forward | 74,847-75,235 | up | *petB* intron | 3 | (47) |
| forward | 75,236-75,649 | up | *petB* intron | 4 | (47) |
| forward | 76,487-77,196 | up | *petD* intron | 5 | (47) |
| forward | 77,740-77,963 | up | *petD* 3' ends; antisense to *petD-rpoA* intergenic | 6 | (37, 47) |
| forward | 77,964-78,112 | up | *petD* 3' ends; antisense to *rpoA* | 7 | (37, 47) |
| forward | 78,113-78,218 | up | *petD* 3' ends; antisense to *rpoA* | 8 | (37, 47) |
| reverse | 71,814-73,668 | up | *psbN* 3' ends; antisense to *psbB* | 9 | NA |
| reverse | 73,669-73,935 | up | *psbN* 3' ends; antisense to *psbB* | 10 | NA |
| reverse | 73,936-74,085 | up | *psbN* 3' ends; antisense to *psbB-psbT* intergenic | 11 | (37) |
| reverse | 74,232-74,365 | down | *psbN* | 12 | (47) |
| reverse | 74,366-75,133 | down | *psbN* 5' ends; antisense to *psbH* and *petB* | 13 | (37) |
| reverse | 75,134-77,383 | up | *rpoA* 3' ends; antisense to *petB* and *petD* | 14 | NA |
| reverse | 77,384-77,605 | up | *rpoA* 3' ends; antisense to *petD* | 15 | NA |

### Benchmarking

#### Data and read mapping

The true positive rate (see *Evaluation metrics*) of DiffSegR and competitors were evaluated on two RNA-Seq datasets comparing *Arabidopsis thaliana* control plants (*col0*) to mutants deficient in the PNPase and Mini-III chloroplast ribonucleases (31, 37). We refer to these datasets as *pnp1-1* and *rnc3/4*, respectively. In the *rnc3/4* dataset both conditions contained two replicates with about 19.5 million reads each while in the *pnp1-1* dataset, both conditions contained two replicates with about 18.6 million reads each. DiffSegR ability to work on a bacterial genome was evaluated using a RNA-Seq dataset comparing a *Bacillus subtilis* control strain (CCB375 strain) to a mutant deficient for the Rae1 ribonuclease (SSB1002 strain) (38). We refer to this dataset as *Δrae1*. Both conditions contained three replicates with about 14.8 million reads each. The IDEAs dataset used to evaluate the false positive rate (see *Evaluation metrics*) contained ten RNA-Seq replicates of the Col-0 *Arabidopsis thaliana* accession grown in nitrogen deficiency condition with about 32.7 million reads each. The plants were grown at the IJPB Phenoscope platform (https://phenoscope.versailles.inrae.fr/) to ensure maximal homogeneity between the replicates (see the GEO database with the accession number GSE234377 for more details). RNA-Seq datasets were aligned to the *Arabidopsis thaliana* chloroplast genome using the OGE pipeline (https://forgemia.inra.fr/GNet/pipelineoge) (39). This pipeline uses the STAR aligner (40). The BAM files corresponding to the aligned *Bacillus subtilis* RNA-Seq dataset were kindly provided by Ciarán Condon. The alignment was performed using the Bowtie aligner (41). These alignments were then used for the downstream analyses. Because DiffSegR is the only evaluated method able to analyze stranded RNA-Seq reads, the BAM files were then split by strand in order to be used by the competing methods and the results for both strands were finally merged.

#### Adjusting method parameters

For the purpose of benchmarking DiffSegR against other methods in terms of true positive rate (see below), one or more parameters likely to change the number and/or the positions of the identified changepoints were adjusted.

1. The minimum depth threshold (*minDepth*) is common to derfinder RL and srnadiff. All contiguous positions with mean of coverages above this threshold are kept. For each method, on both datasets, one hundred *minDepth* values evenly spaced within the interval [1,6000] were tested. The default *minDepth* value of derfinder RL and srnadiff are 5 and 10, respectively.
2. The minimum log2-FC threshold (*minLogFC*) is used by srnadiff to keep only contiguous positions with absolute normalized log2-FC above the threshold. For both methods, on both datasets, one hundred *minLogFC* values evenly spaced within the interval [0.1,7] were tested. The default *minLogFC* value of srnadiff is 0.5.
3. The emission threshold (*emissionThreshold*) is used by srnadiff to define the HMM states. For both methods, on both datasets, one hundred (*emissionThreshold*) values evenly spaced within the interval [0.09, 0.9] were tested. The default *emissionThreshold* value of srnadiff is 0.1.

For all these comparisons and on both datasets, the DiffSegR hyperparameter *λ* was kept to its default value, *λ*=2. In other analyses, all parameters from the different methods tested were set to their default values.

#### Evaluation metrics

At the end of the segmentation process, each method yields a collection of segments that may or may not correspond to genomic regions with differential expression. Differentially expressed regions (DERs) stand for the largest set of segments with a fold-change > 1.5 (symmetrically < 2/3) and a false discovery proportion upper bound set to 5%. Both per-segment fold-change and p-value are estimated using DESeq2 (29). The post-hoc upper bound is obtained by controlling the joint error rate (JER) at significance level of 5% using the Simes family of thresholds implemented in the R package sanssouci (42, 43). Unless specified, all methods were compared using these thresholds. All quality control of the DiffSegR results, including a PCA of counts, a dispersion-mean plot and an histogram of p-values are available in supplementary data for *pnp1-1* (Figures S1-S3), *rnc3/4* (Figures S4-6) and *Δrae1* (Figures S7-S9) datasets. For the comparisons on the *pnp1-1* and *rnc3/4* labeled dataset the error *E* was defined as the total number of labels which are not overlapped by at least one DER. A label is a genomic portion whose corresponding transcript has previously been validated by molecular biology techniques to be differentially accumulated in the mutant compared to WT. The genomic coordinates of the labels can be found in Table S1-S2.

The true positive rate is given by 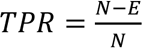 where *N* is the total number of labels. In the blank experiment the replicates of the nitrogen deficiency condition from the IDEAs project were resampled in two groups to test several 2 vs 2, 3 vs 3, 4 vs 4 and 5 vs 5 designs. All the DERs identified are supposed to be false positives. The false positive rate is given by 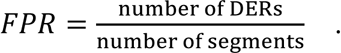

## RESULTS

### Foreword

srnadiff merges the results of a two-level segmentation approach on the per-base p-value (srnadiff HMM) and a three-level segmentation approach on the per-base log2-FC (srnadiff IR) (Figure 1). Consequently, for the purposes of the following comparisons, we will use srnadiff as a representative tool of the methods following similar strategies, including derfinder SB and RNAprof. In addition, due to the lengthy process of estimating the parameters of the model implemented in parseq (days) (20), comparing this tool with srnadiff, derfinder RL and DiffSegR is beyond the scope of our study.

### Speed and memory comparisons

All the simulations presented here were performed with an Intel Core i7-10810U CPU @ 1.10GHz, 16 Go of RAMs and 10 logical cores. On both chloroplast RNA-Seq datasets, DiffSegR returns results in less than 2 minutes. In comparison, it takes less than 30 seconds for a standard differential gene expression (DGE) analysis. The identification of segment boundaries using changepoint detection analysis runs in less than a second on both datasets. The slowest step of the DiffSegR pipeline is the construction of the coverage profiles followed later by the segment count table using the featureCounts program and the BAMs files (Table S3). Less than 1 Go of RAM is necessary and the peak of memory used is reached at the differential analysis step (Table S4).

### DiffSegR facilitates the visualization of DERs

DiffSegR was applied to a RNA-Seq dataset comparing control plants (*col0*) to a mutant deficient in the PNPase chloroplast ribonuclease (*pnp1-1*), a major 3’ processing enzyme (37). When dealing with a gene dense genome like the plastome, annotating a DER using the nearest gene can lead to ambiguities. In this case, visualization of the DERs in a genome viewer, as exemplified for the *psbB-psbT-psbN-psbH-petB-petD* gene cluster (Figure 3), is often the simplest solution. In line with previous molecular studies, DiffSegR identifies 15 DERs, 8 on the forward and 7 on the reverse strand, respectively. For example, the overexpressed segment, in 5’ of the *psbB* gene (DER 1 with genomic positions 72,233 to 72,395) matches an area previously shown to over accumulate RNA 5’ ends in *pnp1-1* (45) and the segment 2 overlapping *psbH* and antisense to *psbN* (DER 2 with genomic positions 74,224 to 74,846) corresponds to various 400 to 700 nt long RNA isoforms previously characterized in *WT* or *pnp1-1* mutants (37, 46–48). The published molecular validations corresponding to the DERs identified in the *psbB* gene cluster by DiffSegR are summarized in Table 1.

### DiffSegR improves the search for DERs

The ability of DiffSegR and competitor methods derfinder and srnadiff (19, 22) to identify DERs was evaluated on two RNA-Seq datasets generated for plants lacking the chloroplast ribonucleases PNPase (see above) and Mini-III (*rnc3/4*) (31, 37). In comparison to control plants, these two mutants over accumulate RNA fragments that are mainly located outside of the annotated genic areas and the RNA-Seq data have been extensively validated using molecular techniques (31, 32). These validations were used to define 23 labels (17 in *pnp1-1* and 6 in *rnc3/4*) where a DER was expected to be found (list and coordinates of the labels in Table S1-S2). Using its default segmentation hyperparameters (λ=2) DiffSegR identified 434 and 25 DERs in the *pnp1-1* and *rnc3/4* datasets respectively (Tables S5-S6; Figures S10- S30), including all the predefined labels (TPR = 1). By contrast, srnadiff and derfinder RL identified 16 and 4 labels out of 17 in *pnp1-1* and 4 and 0 labels out of 6 in *rnc3/4* (Table 2). After adjusting the per-base log2-FC threshold, only srnadiff was also able to reach a TPR of 1 (Figure S31-S34). As expected, standard differential gene expression (DGE) analysis, which relies on known gene annotations and is considered as a routine research tool (3), was unable to identify labels located outside of these annotations, therefore resulting in an TPR of 0. Because the large number of DERs found by DiffSegR could suggest it has a high FPR, we evaluated and compared it to classical DGE analysis (49) using a RNA-Seq dataset containing 10 replicates of the nitrogen deficiency condition. Any DER identified between subsamples of the replicates was therefore considered a false positive. The empirical cumulative distribution functions (eCDFs) of the FPR for both DiffSegR and the DGE analysis were similar when using the 5 vs 5 designs. For the 2 vs 2 designs, approximately 90% and 80% of the designs resulted in less than 2.5% of FPR with DiffSegR and traditional DGE respectively (Figure 4). These observations confirm that the FPR is not inflated in the results of DiffSegR (see Figure S35 for 3v3 and 4v4 designs).

**Figure 4:**
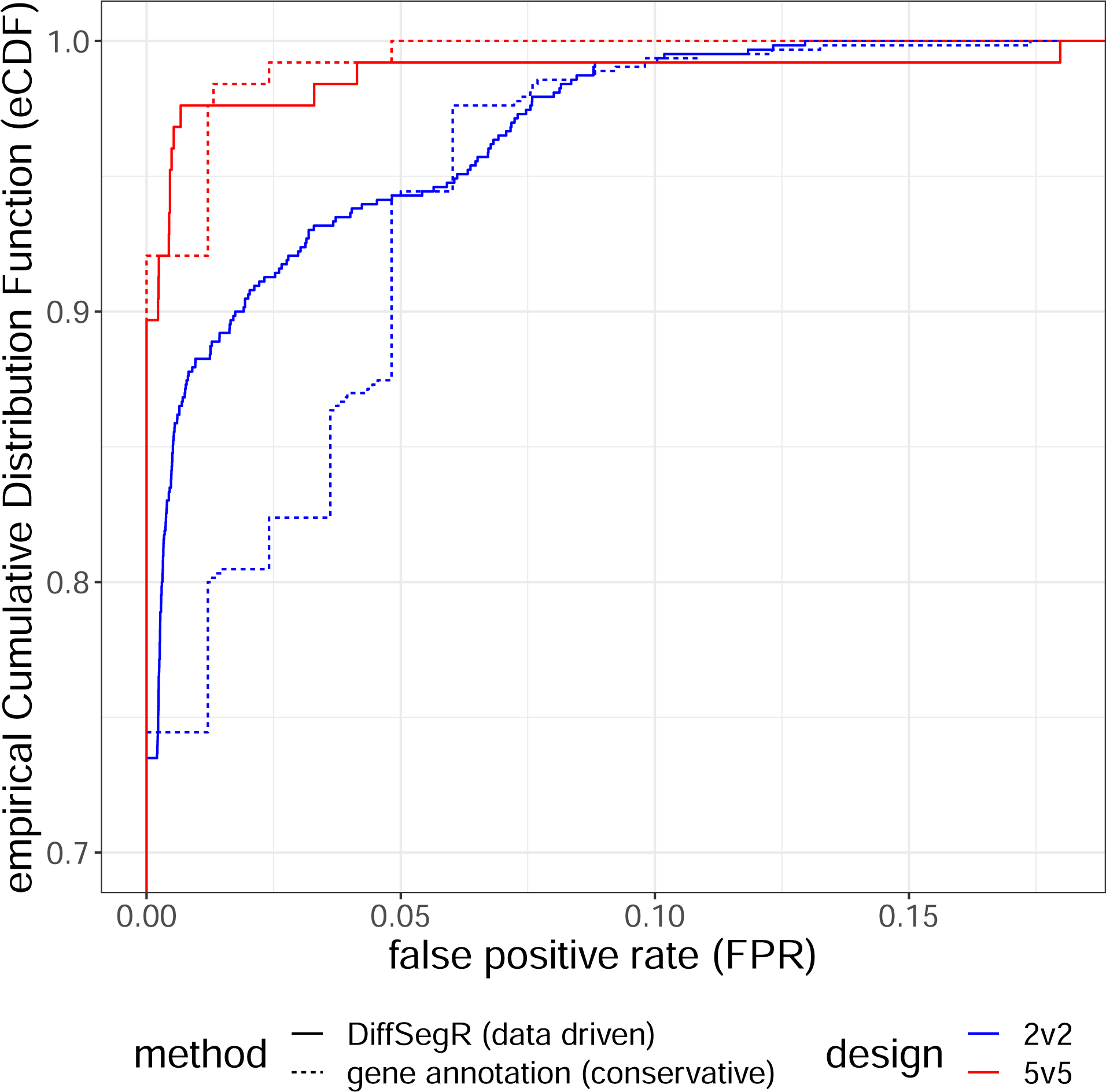
Comparison of the empirical cumulative distribution functions (eCDFs) of the False Positive Rate (FPR) from DiffSegR and the Differential Expression analysis within Gene annotations (DGE). The eCDFs of FPRs from DiffSegR (solid curves) and DGE (dashed curves) methods are compared by re-sampling two groups from 10 biological replicates of the same nitrogen deficiency condition in the IDEAs dataset. The figure displays results for group sizes of 2 (blue curves) and 5 (red curves) (see Figure S35 for 3v3 and 4v4 designs). The eCDF represents the proportion of comparisons (y-axis) with fewer false positives than a specified percentage (x-axis). The eCDF analysis demonstrates that the FPR in DiffSegR results is not inflated compared to the widely-used DGE approach.

**Table 2:** Comparison of the true positive rates (TPRs) for DiffSegR, srnadiff, and derfinder RL methods on the *pnp1-1* (17 labels) and *rnc3/4* (6 labels) datasets. Each method is executed using its default segmentation hyperparameters.

| dataset | method | TPR |
| --- | --- | --- |
| *pnp1-1* | DiffSegR | 1 (17/17) |
| *pnp1-1* | srnadiff | 0.94 (16/17) |
| *pnp1-1* | derfinder RL | 0.24 (4/17) |
| *rnc3/4* | DiffSegR | 1 (6/6) |
| *rnc3/4* | srnadiff | 0.67 (4/6) |
| *rnc3/4* | derfinder RL | 0 (0/6) |

### DiffSegR better captures the differential landscape

Because derfinder RL and srnadiff use a two- or three-level segmentation model they are susceptible to merge in a single DER various contiguous segments having different log2-FC. As a consequence, distinct DER events stemming from distinct RNA maturation processes could be wrongly associated together (Note S4). In contrast, DiffsegR segments the mean of the per-base log2-FC without making any assumption on the number of levels. It should therefore be able to distinguish between contiguous DER events, leading to shorter DER than the other methods. We therefore compared the length distribution of DERs identified by DiffSegR, srnadiff and derfinder RL. In agreement with our expectation, the DERs identified by DiffSegR are on average smaller than those identified by its competitors in both the *pnp1- 1* and *rnc3/4* datasets (Figure 5). Respective median sizes are equal to 211 and 455 nt for DiffSegR and srnadiff (p-value < 2.2*10^-16^, Mann–Whitney U test) in *pnp1-1*. In *rnc3/4* respective median lengths are equal to 15 and 97 nt (p-value = 0.0362, Mann–Whitney U test) (Figure 5.A). An identical trend can be observed between DiffSegR and derfinder RL. In *pnp1-1* respective median sizes are equal to 220 and 826 nt (p-value < 2.2*10^-16^, Mann– Whitney U test). In *rnc3/4*, derfinder fails to detect DERs, accounting for the absence of overlapping DERs between DiffSegR and derfinder RL in this particular dataset (Figure 5.B). We conclude that srnadiff and derfinder RL indeed merge neighboring DERs with different log2-FC.

**Figure 5:**
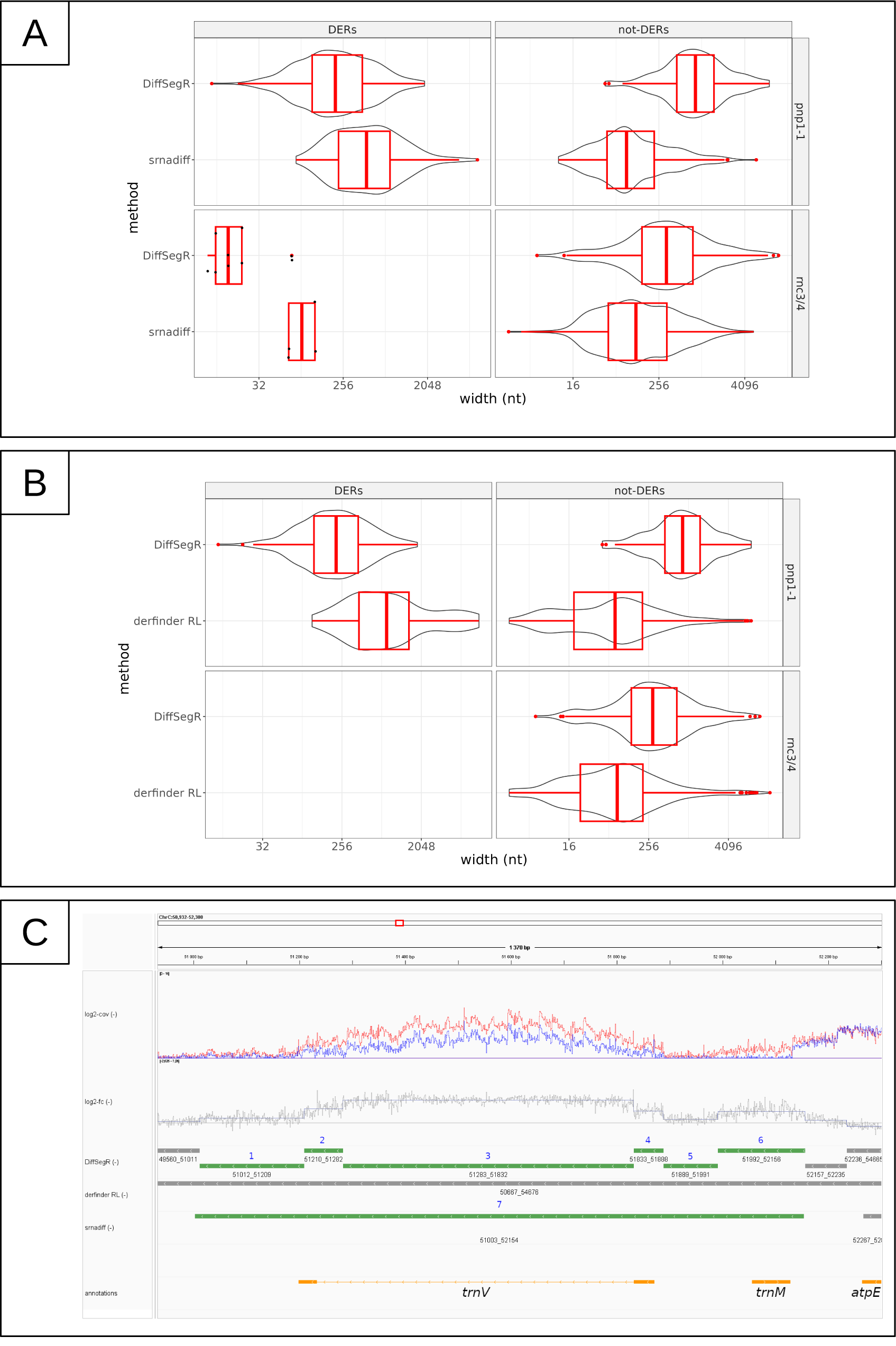
Comparisons of DERs and not-DERs lengths between DiffSegR, derfinder RL and srnadiff on *pnp1-1* and *rnc3/4* datasets. **(A)** The length distribution of DERs and not-DERs identified by DiffSegR and srnadiff are shown using both boxplot and violin plot. Only overlapping (not-)DERs between the compared methods are kept. A (not-)DER of method DiffSegR is considered overlapping either if it covers 90% of a (not-)DER of srnadiff or if 90% of it is covered by a DER of method srnadiff. When there are fewer than 20 overlapping DERs or not-DERs, the violin plot is replaced by a dot plot. **(B)** Similar comparisons were made between DiffSegR and derfinder RL methods. Derfinder does not identify DERs in *rnc3/4*, which explains the lack of overlap between DiffSegR DERs and derfinder RL DERs in this dataset. **(A & B)** In both datasets, DiffSegR not-DERs are on average longer than srnadiff not-DERs and derfinder RL not-DERs in both datasets. Additionally, DiffSegR DERs are on average smaller compared to srnadiff DERs and derfinder RL DERs (Mann-Whitney U test). **(C)** Comparison of DiffSegR, derfinder RL, and srnadiff analyses for the *trnV* gene and the 3’ ends of *atpE*, located on the reverse strand of the chloroplast genome. The tracks are defined as depicted in Figure 3, and further enhanced by incorporating the results from the derfinder RL and srnadiff analysis. DiffSegR identifies 6 up-regulated DERs (IDs 1 to 6). derfinder RL fails to detect any DERs within this region. Lastly, srnadiff discovers a singular DER (ID 7).

Moreover, derfinder RL directly segments the mean of coverages and is therefore susceptible to split regions that are not differentially expressed into distinct segments (Note S5). This is because the shape of the transcriptional signal is strongly influenced by numerous biological and technical factors that are not directly related to *bona fide* transcriptional differences (50). In contrast, DiffSegR uses the per-base log2-FC that is largely unaffected by the underlying transcriptional coverage. This is because local variations in coverage are reproducible and cancel out when taking the difference of the log2 (log2-FC) (Figure S36). As a consequence, we expect DiffSegR to return not-DER longer than derfinder RL. We therefore compared the length distribution of not-DERs identified by DiffSegR, srnadiff and derfinder RL in both *pnp1-1* and *rnc3/4* datasets. Figure 5 shows that the not-DERs identified by DiffSegR are indeed on average longer than those identified by its competitors. Respective median sizes are equal to 833 and 80 nt for DiffSegR and srnadiff (p-value < 2.2*10^-16^, Mann–Whitney U test) in *pnp1-1*. In *rnc3/4* respective median lengths are equal to 294 and 86 nt (p-value < 2.2*10^-16^, Mann–Whitney U test) (Figure 5.A). An identical trend can be observed between DiffSegR and derfinder RL. In *pnp1-1* respective median sizes are equal to 833 and 80 nt (p- value < 2.2*10^-16^, Mann–Whitney U test). In the *rnc3/4* dataset, respective median lengths are equal to 327 and 122 nt (p-value < 2.2*10^-16^, Mann–Whitney U test) (Figure 5.B). We conclude that both srnadiff and derfinder RL over-segment regions that are not differentially expressed in comparison to DiffSegR.

### DiffSegR can be used on sparser genomes

Sparsity refers to the fraction of a genomic region with a null RNA-Seq coverage and is known to cause artifacts in statistical analyses (51). Because the two plant chloroplasts RNA-Seq datasets previously used have a low sparsity ranging from 0.42 to 0.57 we tested DiffSegR on a *Bacillus subtilis* RNA-Seq dataset previously used to decipher the role of the Rae1 ribonuclease (38) and whose sparsity ranged from 0.79 to 0.82 between the different replicates. Using standard differential expression analysis, Leroy et al. identified 46 mRNAs and ncRNAs as significantly up-regulated in the *rae1* mutant (q-value < 0.05 & fold-change > 1.5) and eventually selected seven of them (*S1025*, *S1024*, *S1026*, *yrzI*, *bmrC*, *bmrD*, *bglC*) as candidates for direct degradation by Rae1. DiffSegR returned significant up-regulated DERs overlapping 45 of the 46 genes identified by Leroy et al. including the 7 candidates of interests (Figures S37-S39). In addition, DiffSegR returned significantly up-regulated DERs overlapping 60 other genes (Table S7 and S8). A striking feature was however the over- representation of very short DERs. The five most abundant ones were indeed 4 (6.5%), 6 (6.4%), 5 (5.9%), 2 (5.6%) and 8 (5.4%) nt long while the five most abundant ones in the *pnp1-1* dataset were 55 (1.7%), 73 (1.7%), 83 (1.1%), 204 (1.1%), 56 (0.8%) nt long.

## DISCUSSION

### DiffSegR is a straightforward solution to the DERs detection problem

We here introduced DiffSegR, an R package that allows the discovery of transcriptome-wide expression differences between two biological conditions using RNA-Seq data (Figure 2). While standard RNA-Seq differential analyses rely on reference gene annotations and therefore miss potentially meaningful DERs, DiffSegR directly identifies the boundaries of DERs without requiring any annotation. Unlike its competitors, DiffSegR is designed to analyze stranded RNA-Seq reads, therefore allowing the identification of transcriptional differences on both the forward and reverse strands. This is an invaluable asset when considering the pervasiveness of antisense transcripts (52–54). The output generated by DiffSegR can be easily loaded into the Integrative Genomics Viewer (IGV), providing a user-friendly platform for the exploration and interpretation of the results (Figure 3).

Like other methods willing to automatically identify transcription differences along the genome, DiffSegR addresses a well-defined statistical problem known as the multiple changepoints detection or segmentation problem. Among the many algorithmically and statistically well-established methods that have been developed to tackle this problem (55, 56), DiffSegR uses FPOP (28). This method relies on a Gaussian model to detect changes in the mean of a signal. The computation time of FPOP is log-linear in the signal length, making it time efficient (Table S3). FPOP is statistically grounded (33, 57), and has been shown to be effective in numerous simulations (28, 55) and genomic applications (26, 27, 58). Another advantage of FPOP is that it only has one parameter (the penalty), therefore simplifying calibration and interpretation.

A key feature of DiffSegR is the use of the per-base log2-FC signal for segmentation analysis, a strategy that carries three main advantages. First, it scales with the intensity of the difference up to a normalization constant. Second, it discriminates between up-regulated and down-regulated DERs and third, it is largely insensitive to local variations in coverage as they are reproducible (Figure S36) and cancel out when taking the difference of the logs (log2-FC). Moreover, in contrast to the two-level (DER and not-DER or expressed and not-expressed) and three-level (up-regulated DER, down-regulated DER, not-DER) segmentation models used by other approaches (Figure 1), FPOP does not make any assumptions on the number of levels in the log2-FC and can effectively distinguish between adjacent DERs that involves distinct RNA maturation processes. As a consequence DiffSegR detects fewer changes in non-differential regions but detects more segments in DERs than its competitors (Figure 5). This suggests that DiffSegR is able to effectively summarize the data, providing a detailed and accurate representation of the differential landscape while being more selective in its analysis of not-DERs.

### DiffSegR accurately captures the differential landscape

DiffSegR finds all the extended 3’ and 5’ ends of transcripts, as well as accumulated antisense RNA, in RNA-Seq labeled datasets *pnp1-1* and *rnc3/4*. These labels were previously verified through molecular techniques, and DiffSegR was able to identify them with its default settings, while none of the competitors tested were able to do so. However, the use of the same dataset twice in DiffSegR (and its competitors), a procedure so-called double dipping, first for segmentation and then for differential analysis may result in an inflated false positive rate (59–61). We therefore verified that the FPR of DiffSegR is similar to standard DGE analysis using a blank experiment (Figure 4). A possible explanation to the observed robustness is the fact that DiffSegR uses different aspects of the data in its two steps: while the segmentation uses the per-base log2-FC, the DEA relies on normalized counts, per-segment log2-FC, and dispersion. The last three parameters are estimated by DESeq2.

We are therefore confident that the numerous DERs identified outside of the predefined labels in the two chloroplastic RNA-Seq datasets represent *bona fide* DERs. For example, 387 out of the 434 DERs identified in the *pnp1-1* RNA-Seq experiment did not overlap labels. While an exhaustive molecular validation of these 387 segments is beyond the scope of this study, numerous evidences suggest they are accurate. Specifically, DiffSegR identifies 72 DERs overlapping all the 25 plastid introns in the PNPase mutant, a feature previously shown to reflect a lack of intron degradation following splicing in the mutant (47). Neither srnadiff nor derfinder RL were able to capture this feature entirely. Another example suggesting that DiffSegR does not over-segment the differential transcription profile is displayed for genomic area 51,012-52,156 in Figure 5.C. While it is not differentially expressed according to derfinder RL, srnadiff considers it as a single DER (DER 7 with genomic positions 51,003 to 52,154) and DiffSegR identifies 6 contiguous different DERs within it. The multiplicity of DERs identified by DiffSegR seems to better reflect the shape of the log2-FC and is also consistent with the known roles of the PNPase in transcript 3’ end maturation (DER 1 with genomic positions 51,012 to 51,209 and DER 6 with genomic positions 51,992 to 52,156 for *trnV* and *atpE*, respectively) or the degradation of tRNA 5’ precursor (DER 5 with genomic positions 51,889 to 51,991 for *trnV*) (32). Finally, both *trnV* exons over accumulate (DERs 2 and 4 with genomic positions 51,210 to 51,282 and 51,833 to 51,888, respectively) in the mutant, along with the corresponding intron (DER 3 with genomic positions 51,283 to 51,832). The segmentation in three different DERs is, once again, an accurate interpretation of the two different biological mechanisms targeting tRNAs and introns in the mutant (47, 62).

### Larger genomes with more zeroes

DiffSegR is also effective and powerful on genomes larger and more complex than the chloroplast. It effectively identified the two RNA locations that have been shown to be degraded by the Rae1 endoribonuclease in *Bacillus subtilis* (38, 63)(Deves et al. 2023; Leroy et al. 2017). This illustrates one of the big advantages of DiffSegR, it can be easily used to narrow down the number of genomic regions worth investigating. From the 4.2 Gb *Bacillus* genome it identified 1833 regions (Table S7) that contained the two known cleavage sites, a number that is compatible with the workforce of most research teams. It is however true that the segmentation model used by DiffSegR may result in an over segmentation in profiles containing many base pairs with a null coverage. This could be problematic when addressing even larger genomes, like nuclear ones, and prevent interpretability of the results.

A straightforward solution would be to apply DiffSegR to smaller portions of the genome, only keeping the ones with sufficient coverage. This however comes with issues of its own as (i) identifying those genomic portions is a segmentation problem itself, multiplying the genomic areas complexifies selection, and (ii) this leads to a *triple-dipping* problem as the data is used three times (identification of the genomic area, segmentation within the genomic area and differential expression analysis). Alternative strategies would be to integrate more advanced segmentation methods already available. More specifically, we believe it could be useful to (i) weight the base pair according to its coverage (using a weighted version of FPOP, (64)), (ii) consider full length reads at the prize of modeling auto-correlation (65), and (iii) model the discrete nature of the data using a negative binomial model (66).

## Conclusion

In conclusion, DiffSegR is a powerful tool that provides researchers with a systematic and accurate way to discover expression differences between two conditions using RNA-Seq data, without the need for prior annotations. Because it is designed to compare two conditions, we believe that DiffSegR has the potential to change the way researchers approach differential expression analysis, especially considering the wealth of RNA-Seq based strategies aimed at capturing specific events (67). For example, it has already been used on RNA immunoprecipitation sequencing data to study translation initiation in plant mitochondria (68). We anticipate it could similarly be used to find newly transcribed RNAs compared to mature RNA control in nascent RNA analysis (69), to find differences in ribosome bound RNA in translatome analysis (70) or to discriminate structured (double-stranded RNA) from unstructured RNAs in structurome analysis (71). We expect that the use of DiffSegR will lead to new discoveries and insights in the field of transcriptomic.

## DATA AVAILABILITY

### Software availability

The latest version of the DIffSegR R package is available at https://aliehrmann.github.io/DiffSegR/index.html. The package includes a Vignette which shows on a minimal example how to use the main functions.

### Data availability

● Raw sequences for the *rnc3/4* dataset have been retrieved from the BioProject database with the accession number PRJNA268035.
● Raw sequences for the *pnp1-1* dataset have been retrieved from the SRA database with the accession number SRA046998.
● Raw sequences for the nitrogen deficiency condition from the IDEAs dataset are available at GEO database with the accession number GSE234377.
● Raw sequences for the *Δrae1* dataset can be accessed from the GEO database with the number GSE93894.

### Reproducibility

The scripts used to generate the figures/tables from this manuscript and figures/tables from the Supplementary Materials are available at https://github.com/aLiehrmann/DiffSegR_paper.

## ACKNOWLEDGEMENTS

The authors would like to thank Ciarán Condon for providing the sequencing data about the *B.subtilis Δrae1* mutant and extensive discussions about the analyses. This work has benefited from the support of IJPB’s Plant Observatory technological platforms. We also thank Amber Hotto for help proofreading the manuscript.

## FUNDING

This work was supported by the Agence Nationale de la Recherche through the grant ANR- 20-CE20-0004 JOAQUIN to BC. The IDEAS experiment was funded by an ATIGE grant from Génopole. AL was supported by a PhD fellowship from the French ministère de l’enseignement supérieur et de la recherche. The IPS2 and IJPB benefit from the support of Saclay Plant Sciences-SPS (ANR-17-EUR-0007).

## CONFLICT OF INTEREST

None declared.

## Notes

### Competing Interest Statement

The authors have declared no competing interest.

### Summary of Updates

- We have now numbered the equations in the 'DiffSegR segmentation model' section (equation 1, equation 2 and equation 3). Additionally, we have cited these equations in the relevant paragraphs: namely, the 'Segmentation model' paragraph, 'Estimation of the segments' paragraph, 'Normalization' paragraph, and the 'Overview of the DiffSegR package' section. - We have revised the 'DiffSegR segmentation model' section to distinctly delineate the data, the theoretical model, and the parameter estimation of the model. The 'Differential transcription profile' paragraph describes formally the computation of coverage profiles and then the per-base log2-FC. We then introduce a new paragraph, 'Segmentation model', which elucidates the theoretical model for segmentation. The subsequent paragraph, 'Estimation of the segments', provides a detailed overview of the estimation of changepoints using the FPOP algorithm. - Furthermore, acknowledging the coherence concern, we have moved the 'Data and read mapping' paragraph. While it was initially located in the 'DiffSegR segmentation model' section, it is now located in the 'Benchmarking' section to maintain thematic consistency.

