## Supplementary Materials for "DiffSegR: An RNA-Seq data driven method for differential expression analysis using changepoint detection"

**The supplementary file includes:**

Figures S1 to S39;

Notes S1 to S5 containing the supplementary Table S9 and the supplementary Figures S40 to S44;

### Supplementary Figures S1 to S39

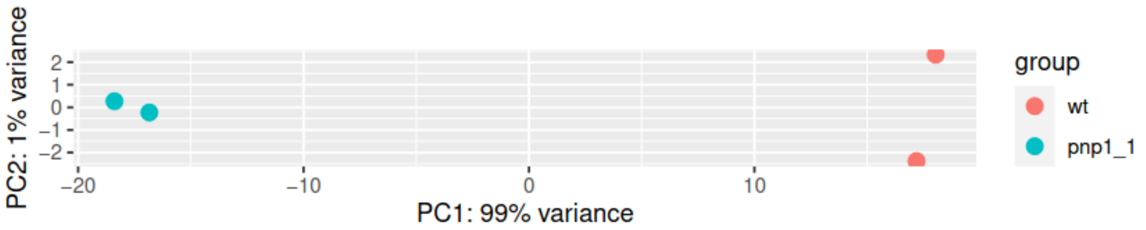


**Figure S1:** PCA of transformed counts from *pnp1-1* RNA-Seq experiment analyzed with DiffSegR. The biological replicates cluster well by condition on PC1.

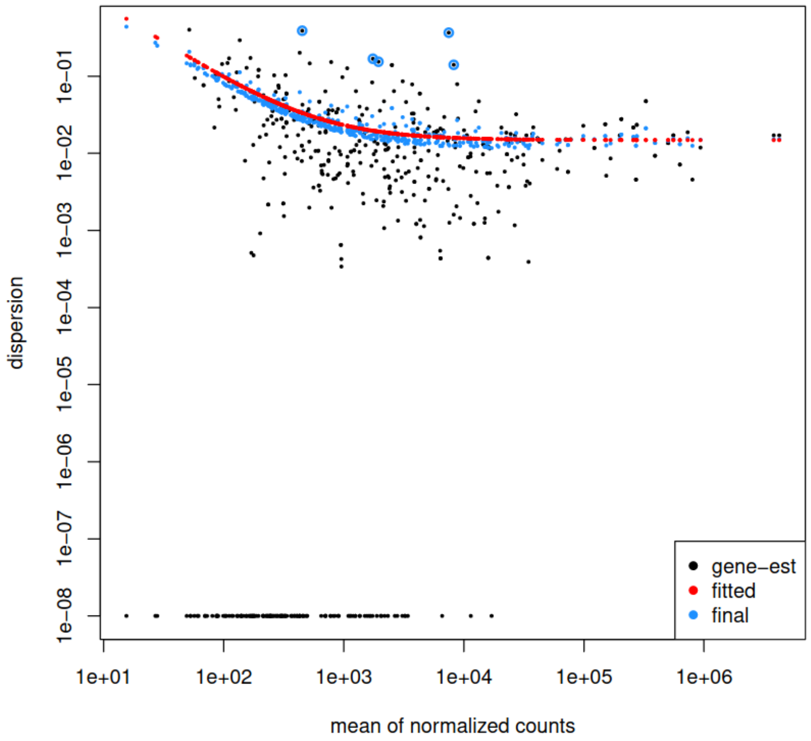


**Figure S2:** Dispersion-mean plot from *pnp1-1* RNA-Seq experiment analyzed with DiffSegR. The plot shows a characteristic dispersion-mean trend for RNA-Seq data.

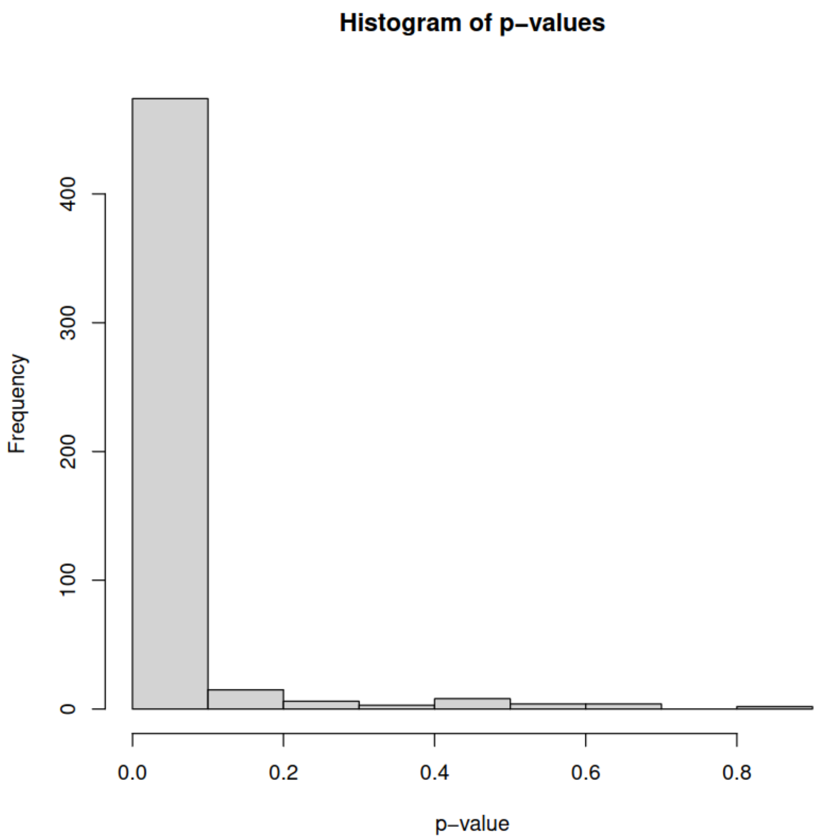


**Figure S3:** Histogram of p-values from *pnp1-1* RNA-Seq experiment analyzed with DiffSegR. The histogram does not show oddity.

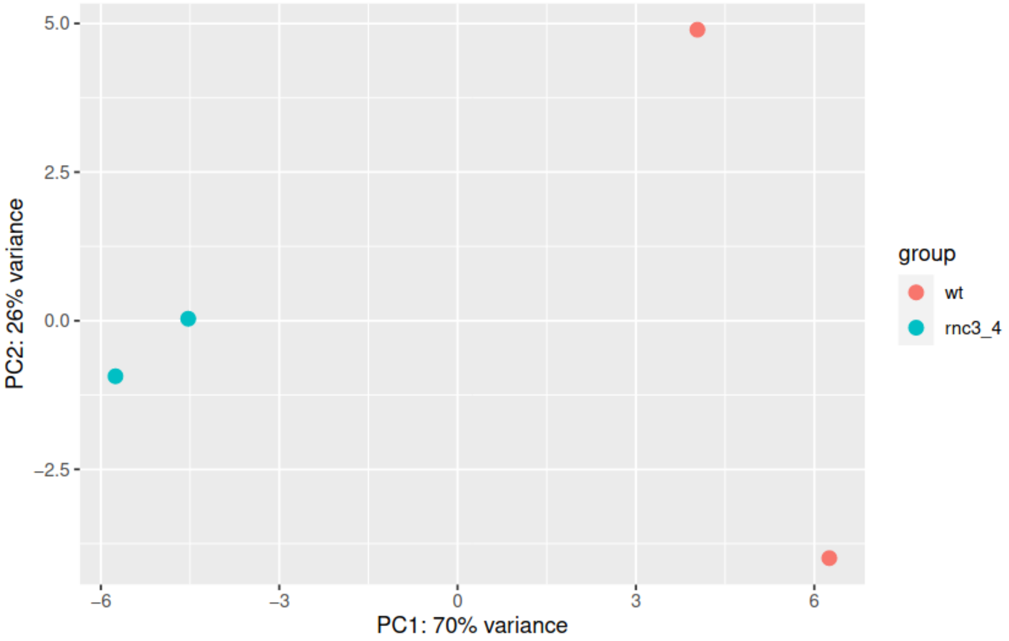


**Figure S4**: PCA of transformed counts from *rnc3/4* RNA-Seq experiment analyzed with DiffSegR. The biological replicates cluster well by condition on PC1.

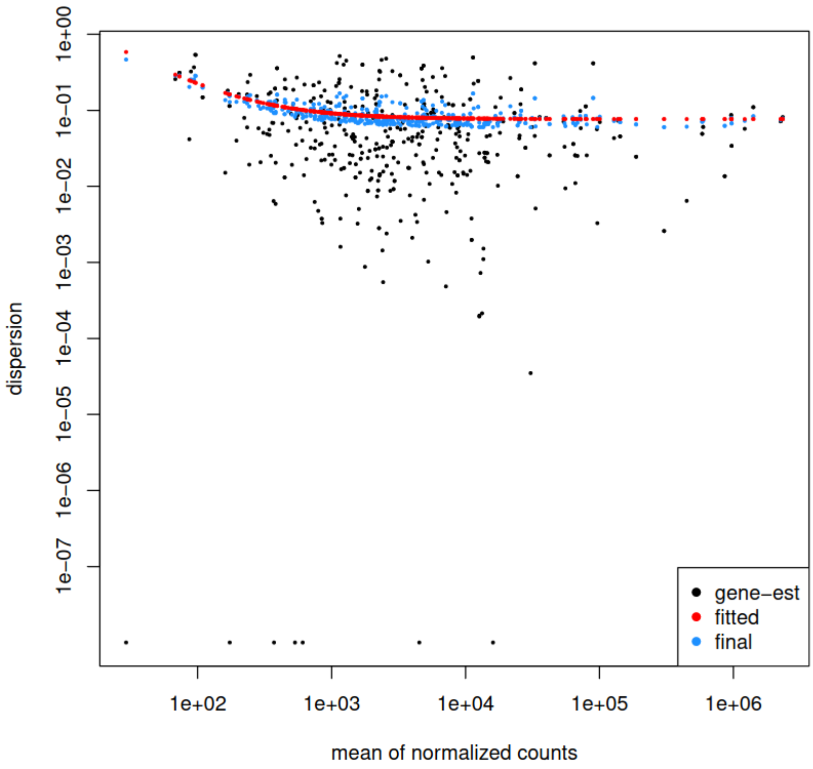


**Figure S5:** Dispersion-mean plot from *rnc3/4* RNA-Seq experiment analyzed with DiffSegR. The plot shows a characteristic dispersion-mean trend for RNA-Seq data.

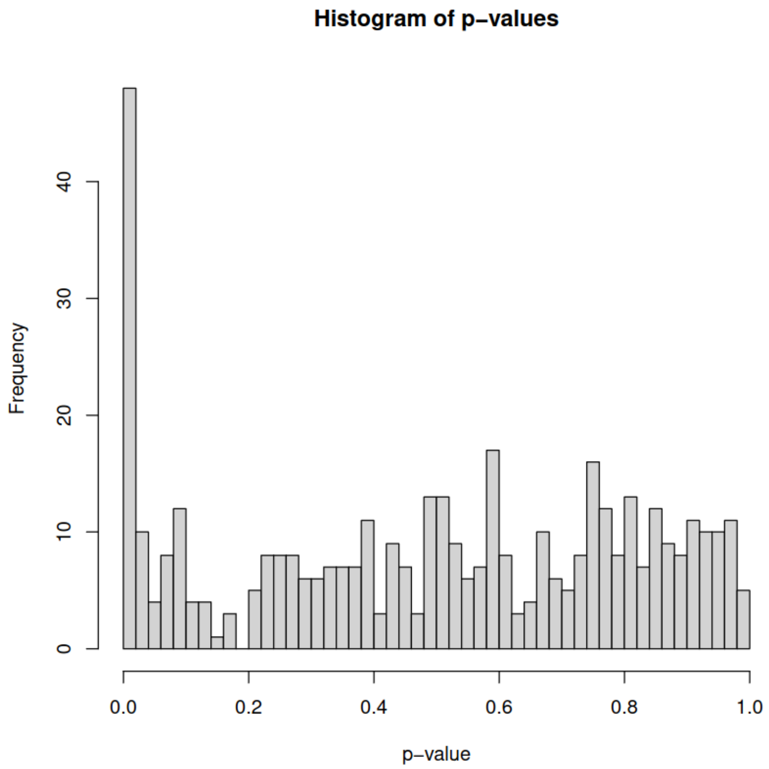


**Figure S6**: Histogram of p-values from *rnc3/4* RNA-Seq experiment analyzed with DiffSegR. The histogram does not show oddity.

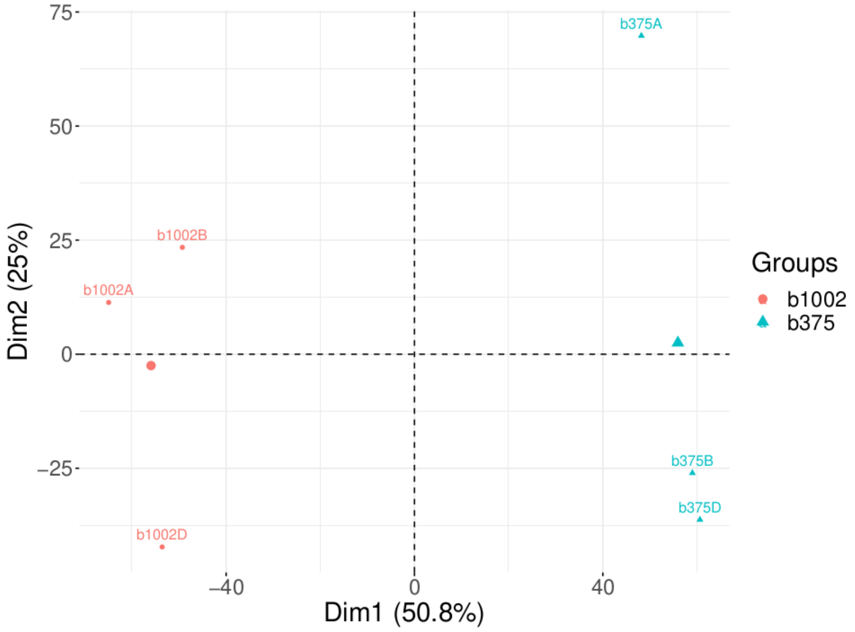


**Figure S7**: PCA of transformed counts from $\Delta$*rae1* RNA-Seq experiment analyzed with DiffSegR. The biological replicates cluster well by condition on Dim1.

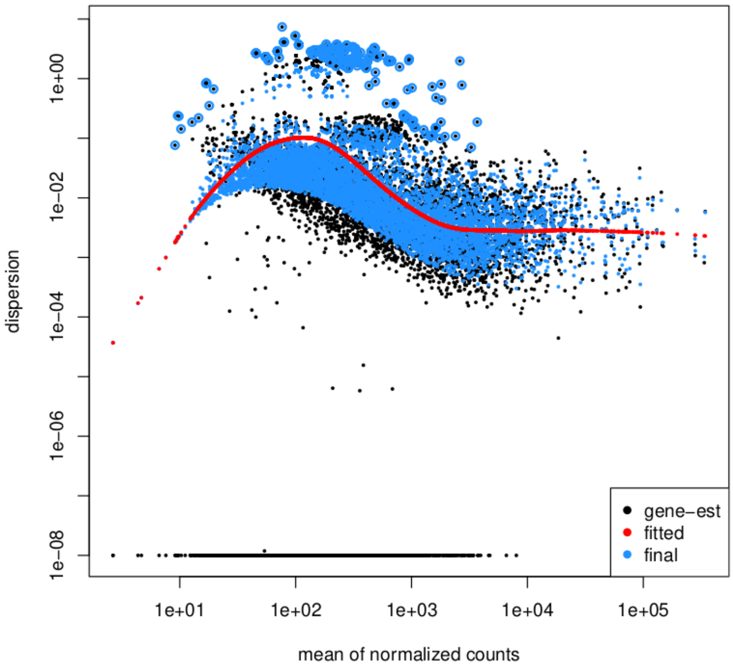


**Figure S8**: Dispersion-mean plot from $\Delta$*rae1* RNA-Seq experiment analyzed with DiffSegR. The dispersion of small counts is negatively biased. Small counts can be filtered out during the preprocessing step.

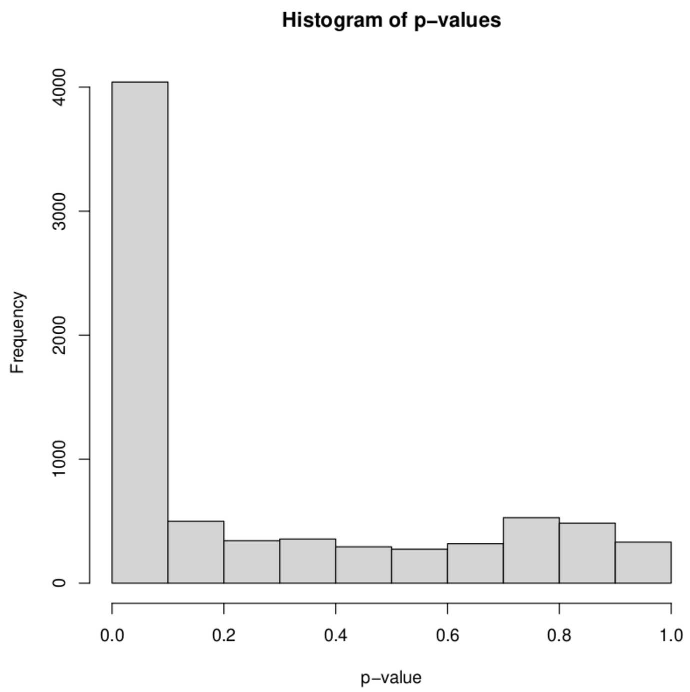


**Figure S9**: Histogram of p-values from $\Delta$*rae1* RNA-Seq experiment analyzed with DiffSegR. The histogram does not show oddity.

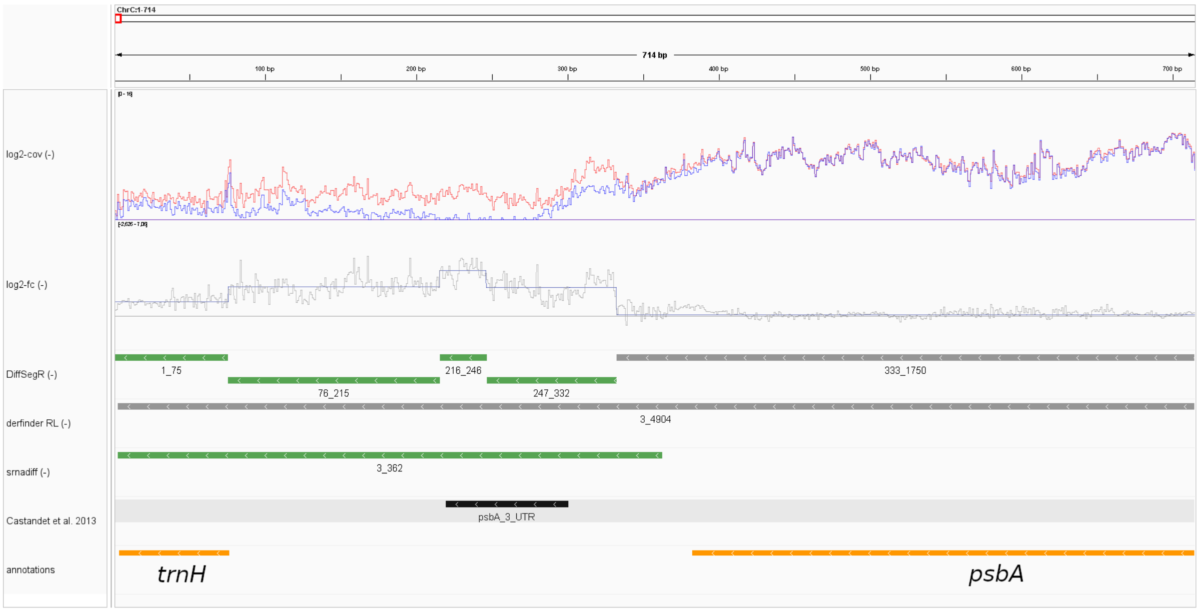


**Figure S10**: Comparison of DiffSegR, derfinder RL, and srnadiff analyses of chloroplast genomic positions 1 to 714 on the reverse strand in the *pnp1-1* dataset. The tracks from top to bottom represent: (log2-Cov (-)) the mean of coverages on the log2 scale for the reverse strand in both biological conditions of interest, with the blue line representing the WT condition and the red line representing the *pnp1-1* condition; (log2-FC (-)) the per-base log2-FC between *pnp1-1* (numerator) and WT (denominator) for the reverse strand. The straight horizontal line represents the zero indicator. When the per-base log2-FC is above or below the zero indicator line, it suggests up-regulation or down-regulation, respectively, in *pnp1-1* compared to WT. The changepoint positions are indicated by vertical blue lines, and the mean of each segment is shown by horizontal blue lines connecting two changepoints; (DiffSegR (-)) the differential expression analysis results for segments identified by DiffSegR on the reverse strand are presented as follows: up-regulated regions are depicted in green, down-regulated regions in purple, and non-differentially expressed regions (non-DERs) in gray. (derfinder RL (-)) the derfinder RL results in the same format as the previous track; (srnadiff (-)) the srnadiff results in the same format as the previous track. (Castandet et al. 2013) the labels of differentially accumulated RNAs in *pnp1-1* compared to WT based on molecular biology validations described in Castandet et al. 2013; (annotations) the genes annotations. The bedGraph and gff3 files used to generate the tracks and the xml file used to load them in IGV were created using the *exportResults* function of the DiffSegR R package. The session was loaded in IGV 2.12.3 for Linux.

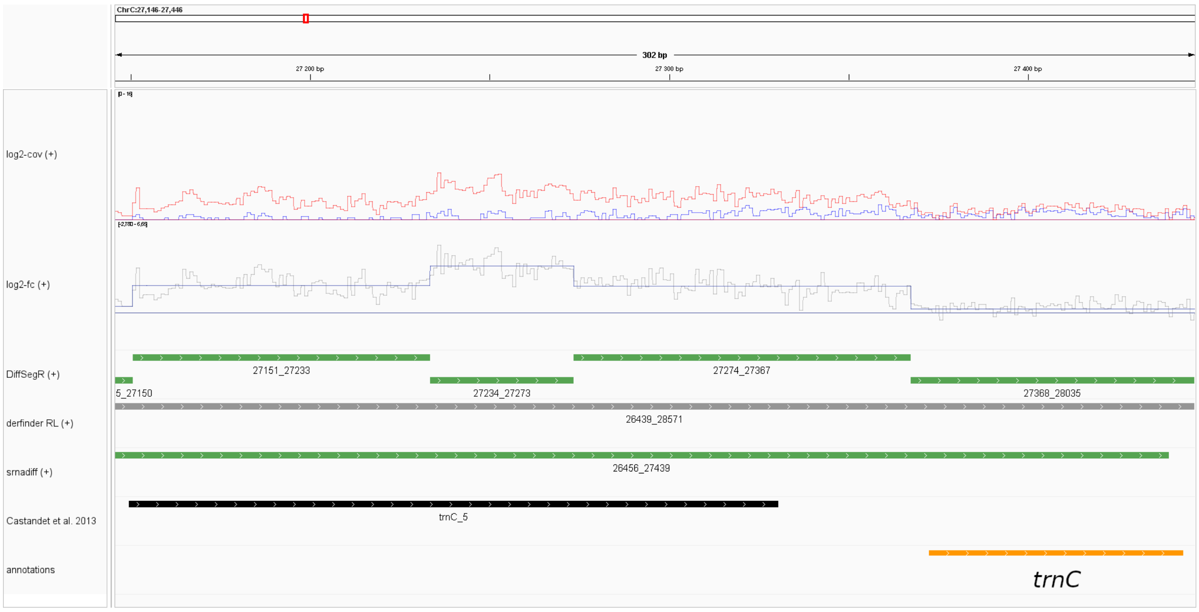


**Figure S11:** Comparison of DiffSegR, derfinder RL, and srnadiff analyses of chloroplast genomic positions 27,146 to 27,446 on the forward strand in the *pnp1-1* dataset. The tracks are similar to those described in Figure S10.

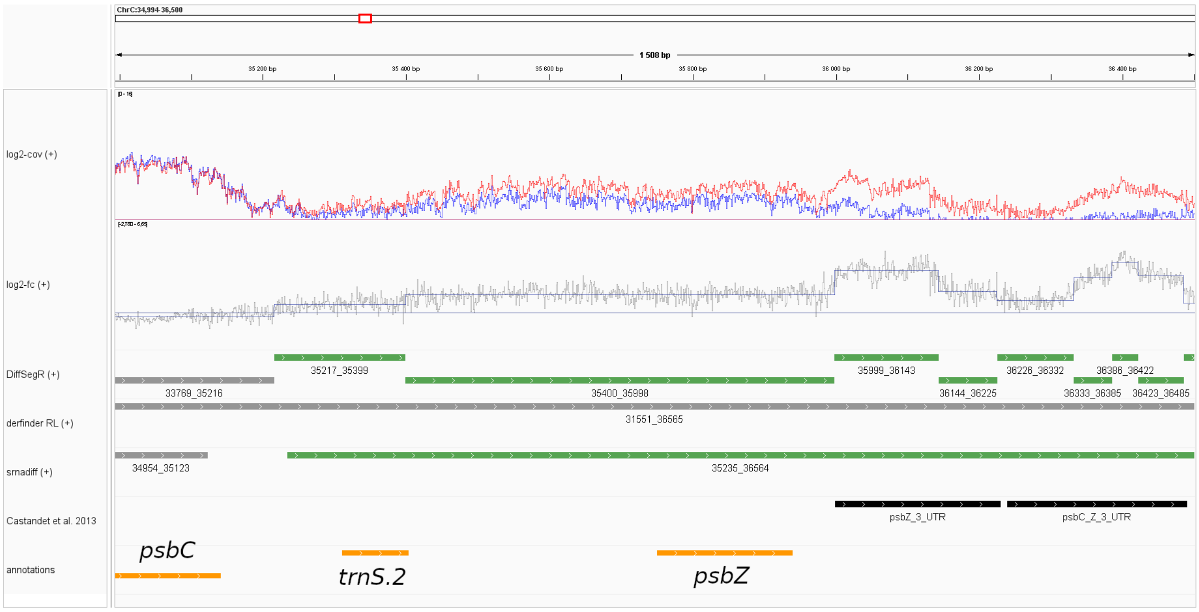


**Figure S12**: Comparison of DiffSegR, derfinder RL, and srnadiff analyses of chloroplast genomic positions 34,994 to 36,500 on the forward strand in the *pnp1-1* dataset. The tracks are similar to those described in Figure S10.

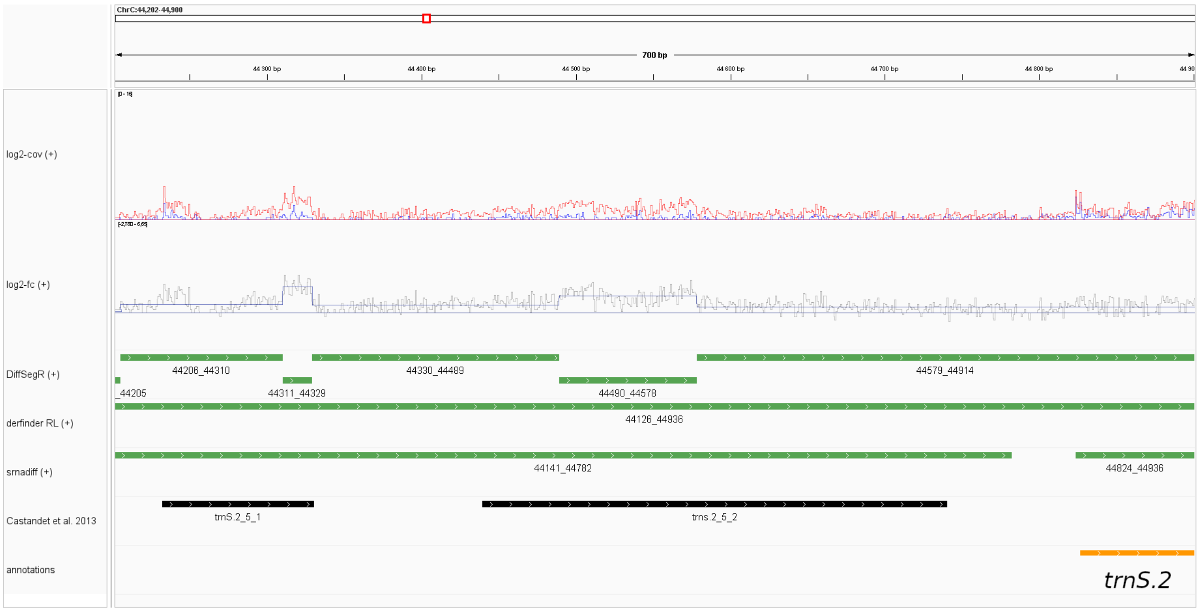


**Figure S13:** Comparison of DiffSegR, derfinder RL, and srnadiff analyses of chloroplast genomic positions 44,202 to 44,900 on the forward strand in the *pnp1-1* dataset. The tracks are similar to those described in Figure S10.

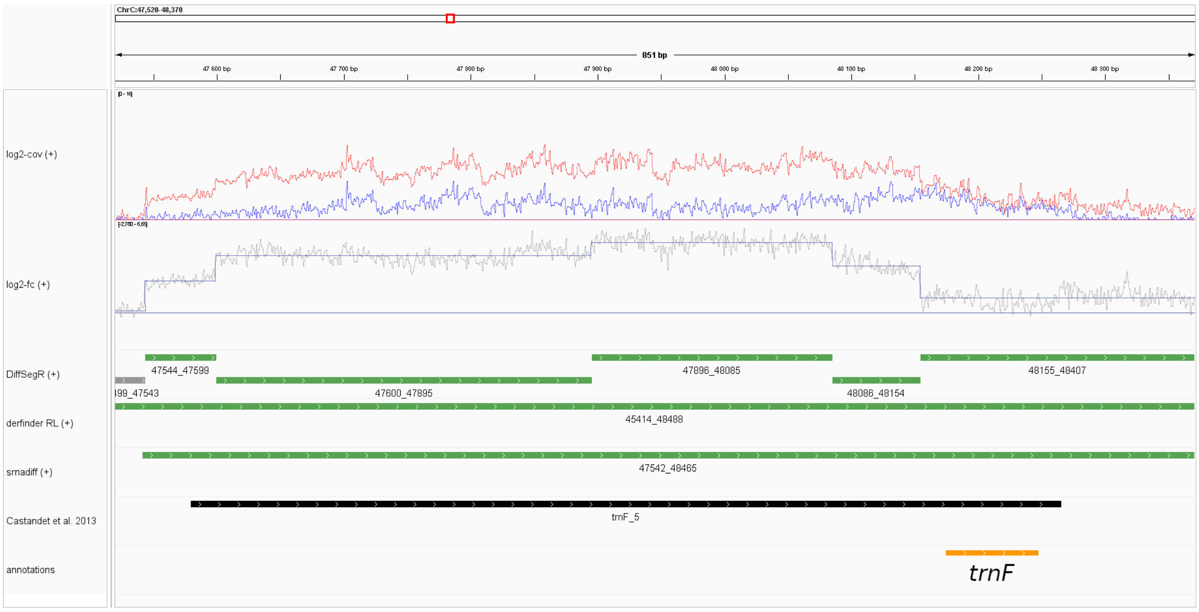


**Figure S14:** Comparison of DiffSegR, derfinder RL, and srnadiff analyses of chloroplast genomic positions 47,520 to 48,370 on the forward strand in the *pnp1-1* dataset. The tracks are similar to those described in Figure S10.

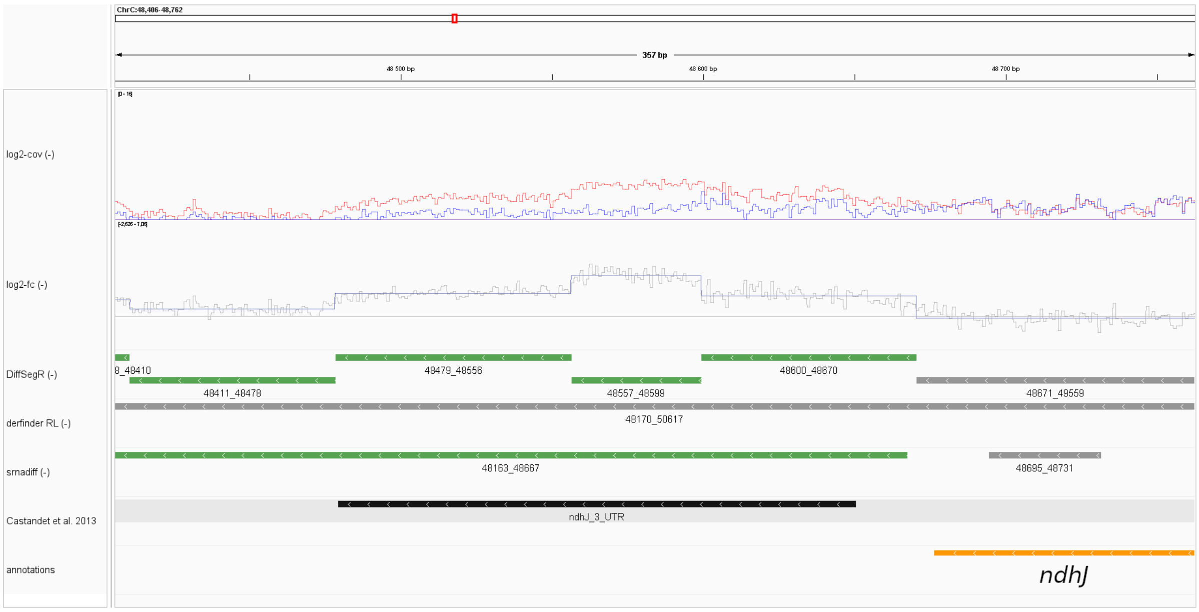


**Figure S15:** Comparison of DiffSegR, derfinder RL, and srnadiff analyses of chloroplast genomic positions 48,406 to 48,762 on the reverse strand in the *pnp1-1* dataset. The tracks are similar to those described in Figure S10.

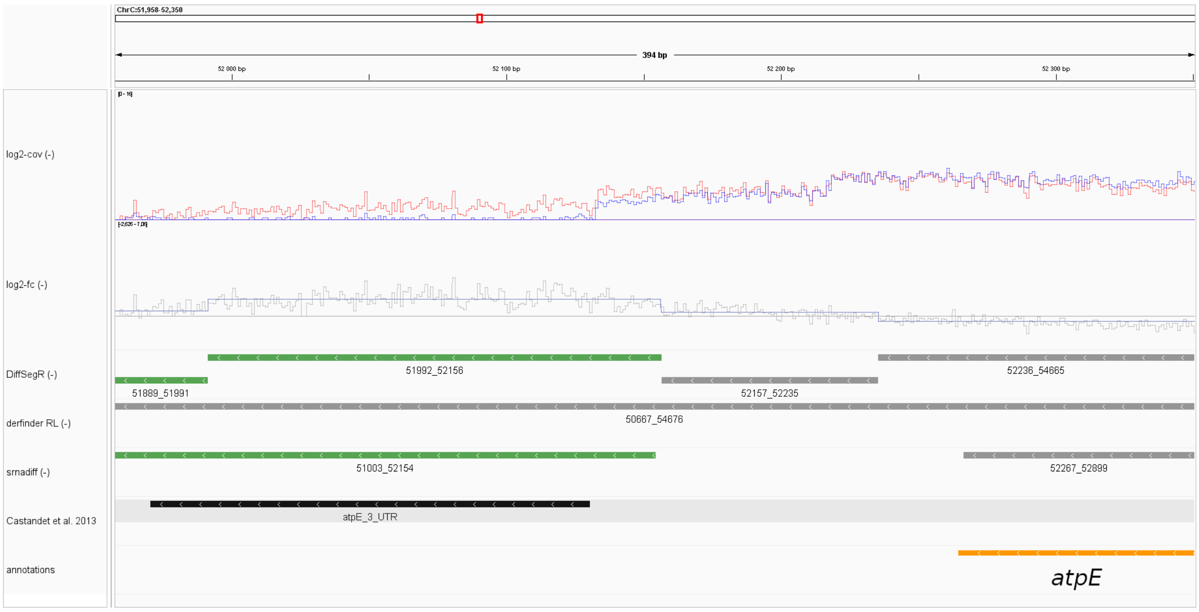


**Figure S16:** Comparison of DiffSegR, derfinder RL, and srnadiff analyses of chloroplast genomic positions 51,958 to 52,350 on the reverse strand in the *pnp1-1* dataset. The tracks are similar to those described in Figure S10.

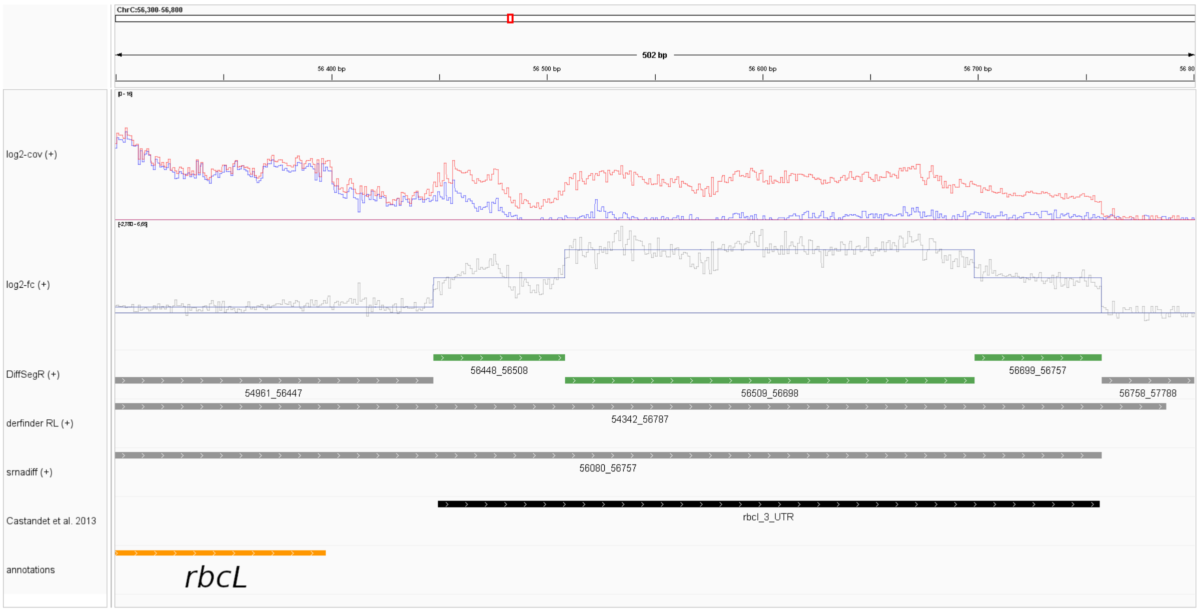


**Figure S17:** Comparison of DiffSegR, derfinder RL, and srnadiff analyses of chloroplast genomic positions 56,300 to 56,800 on the forward strand in the *pnp1-1* dataset. The tracks are similar to those described in Figure S10.

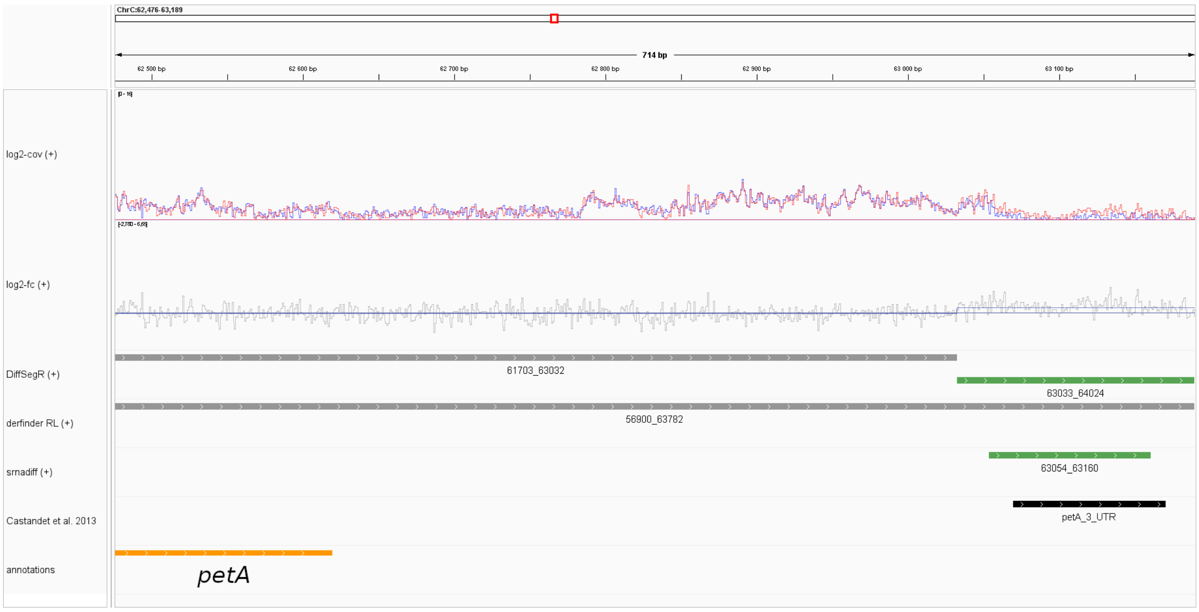


**Figure S18:** Comparison of DiffSegR, derfinder RL, and srnadiff analyses of chloroplast genomic positions 62,476 to 63,189 on the forward strand in the *pnp1-1* dataset. The tracks are similar to those described in Figure S10.

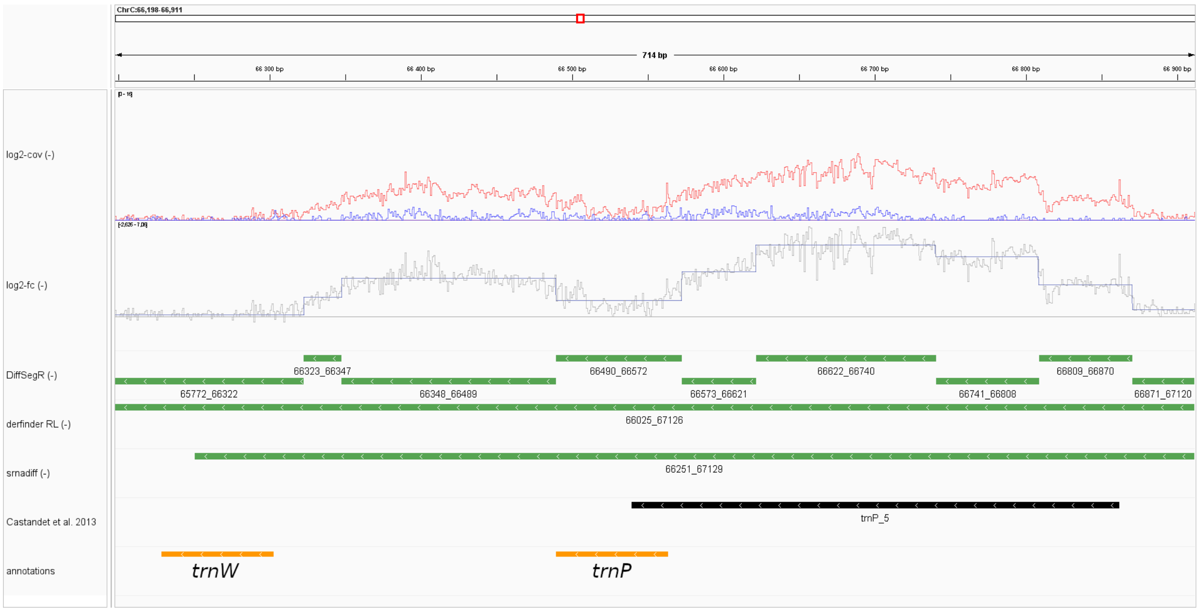


**Figure S19:** Comparison of DiffSegR, derfinder RL, and srnadiff analyses of chloroplast genomic positions 66,198 to 66,911 on the reverse strand in the *pnp1-1* dataset. The tracks are similar to those described in Figure S10.

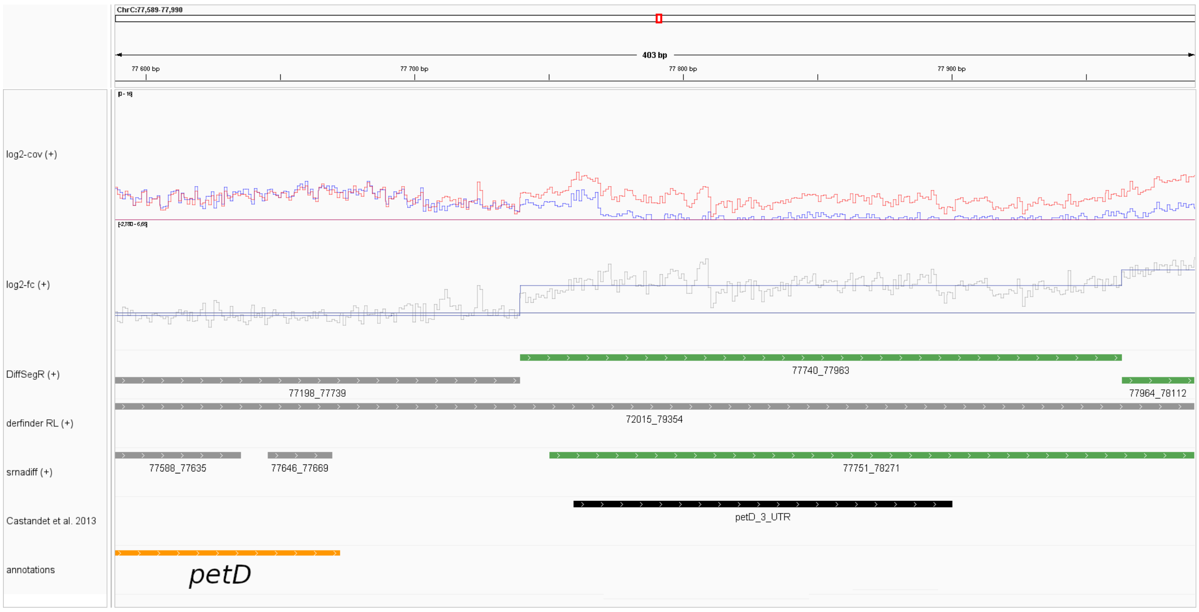


**Figure S20:** Comparison of DiffSegR, derfinder RL, and srnadiff analyses of chloroplast genomic positions 77,589 to 77,990 on the forward strand in the *pnp1-1* dataset. The tracks are similar to those described in Figure S10.

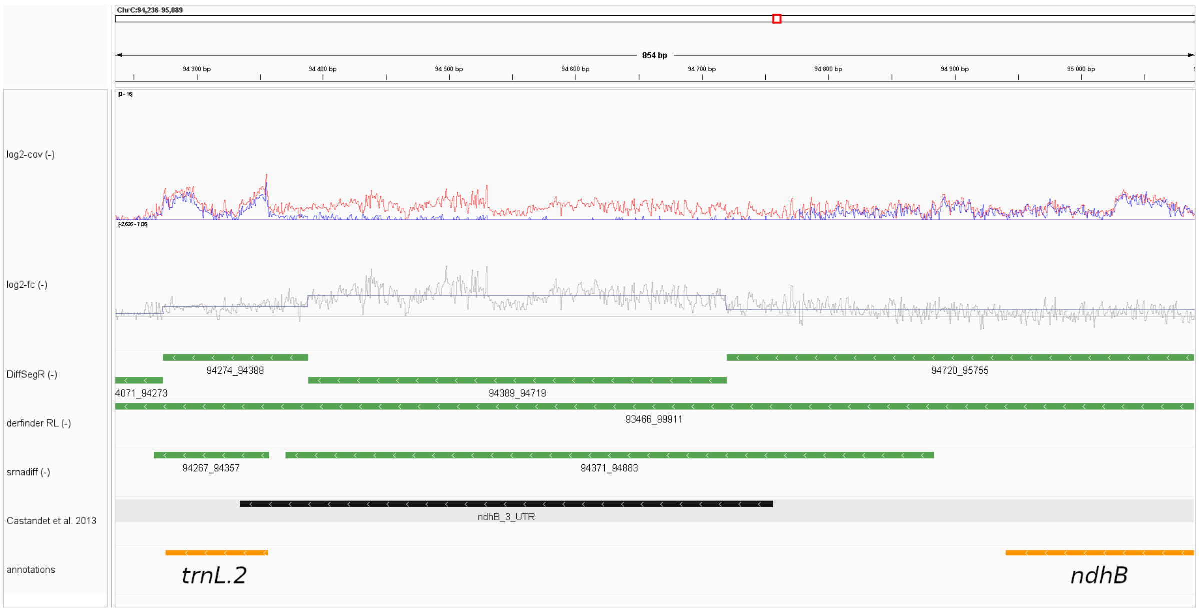


**Figure S21:** Comparison of DiffSegR, derfinder RL, and srnadiff analyses of chloroplast genomic positions 94,236 to 95,089 on the reverse strand in the *pnp1-1* dataset. The tracks are similar to those described in Figure S10.

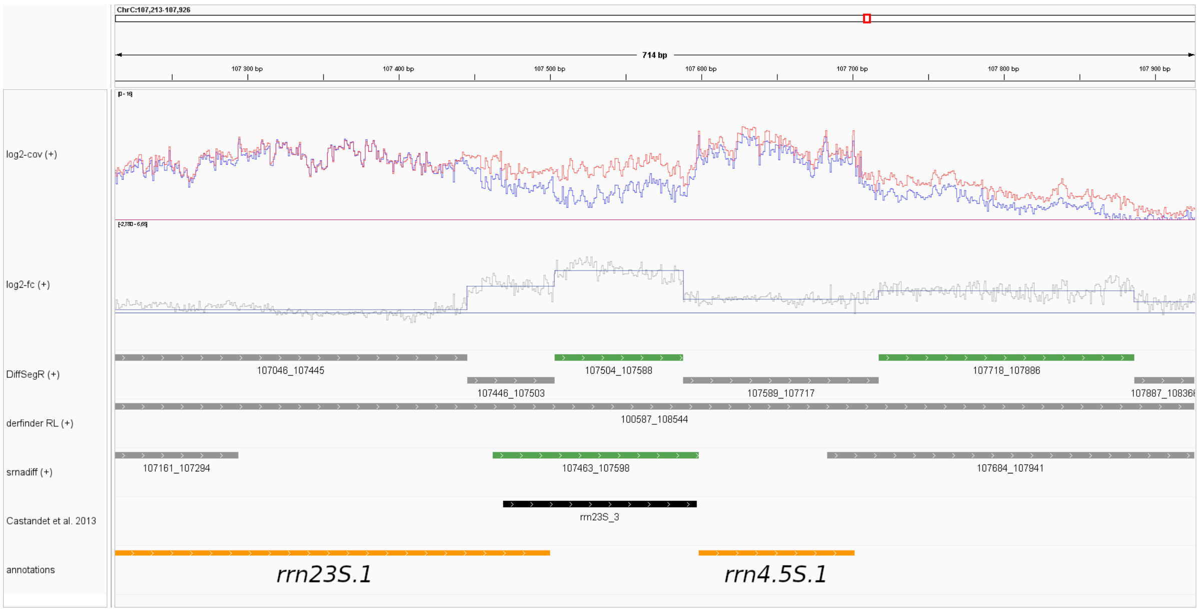


**Figure S22:** Comparison of DiffSegR, derfinder RL, and srnadiff analyses of chloroplast genomic positions 107,213 to 107,926 on the forward strand in the *pnp1-1* dataset. The tracks are similar to those described in Figure S10.

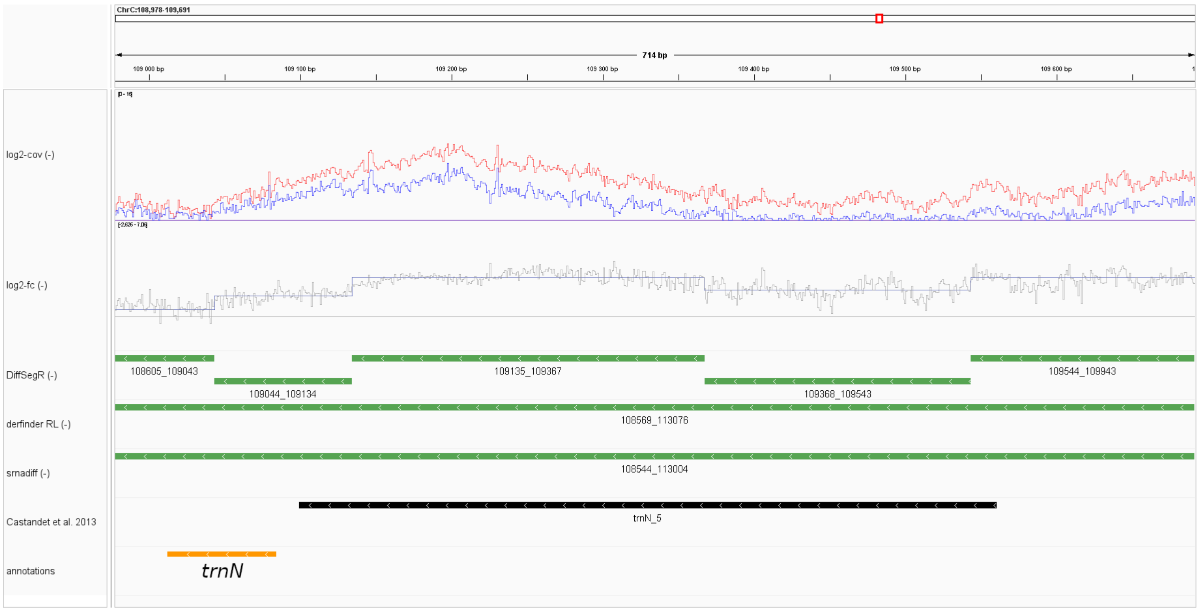


**Figure S23:** Comparison of DiffSegR, derfinder RL, and srnadiff analyses of chloroplast genomic positions 108,978 to 109,691 on the reverse strand in the *pnp1-1* dataset. The tracks are similar to those described in Figure S10.

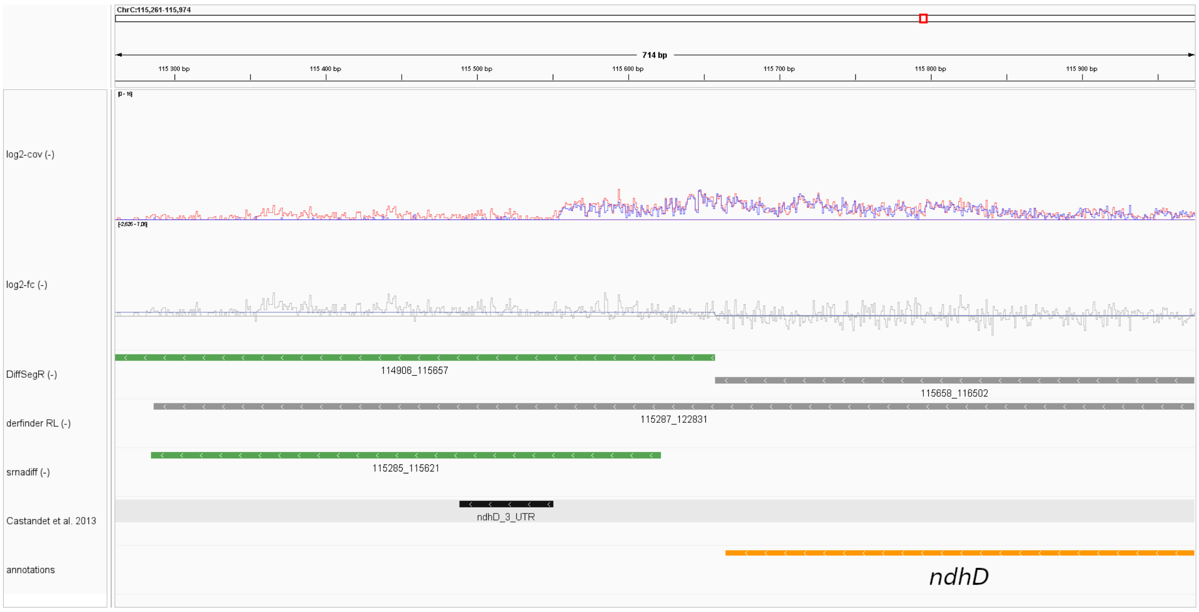


**Figure S24:** Comparison of DiffSegR, derfinder RL, and srnadiff analyses of chloroplast genomic positions 115,261 to 115,974 on the reverse strand in the *pnp1-1* dataset. The tracks are similar to those described in Figure S10.

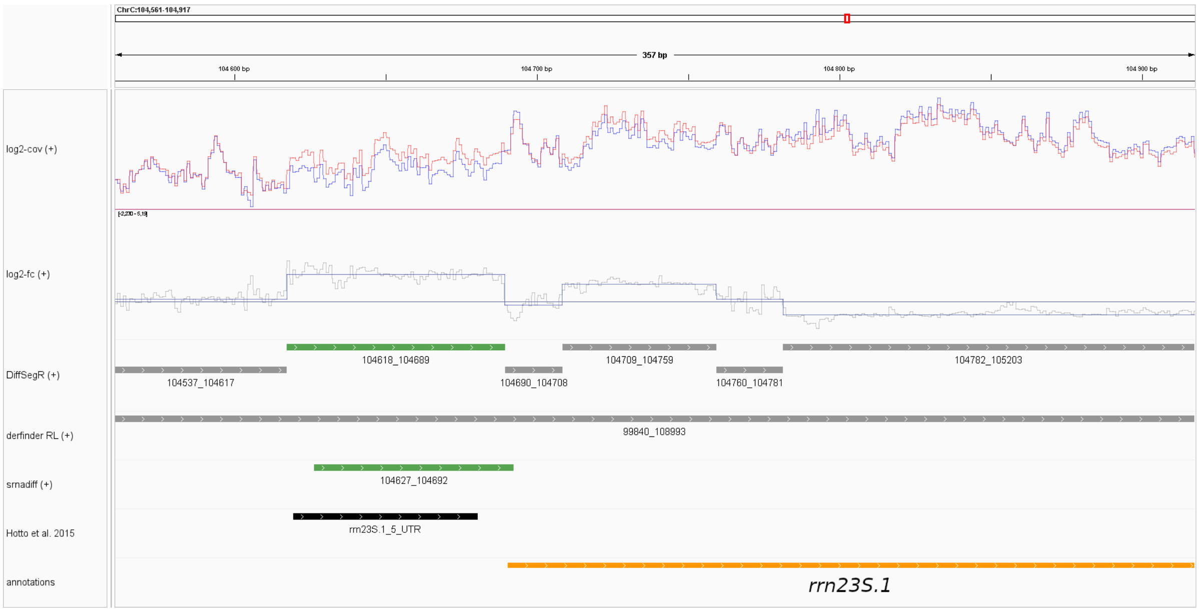


**Figure S25:** Comparison of DiffSegR, derfinder RL, and srnadiff analyses of chloroplast genomic positions 104,561 to 104,917 on the forward strand in the *rnc3/4* dataset. The tracks are similar to those described in Figure S10.

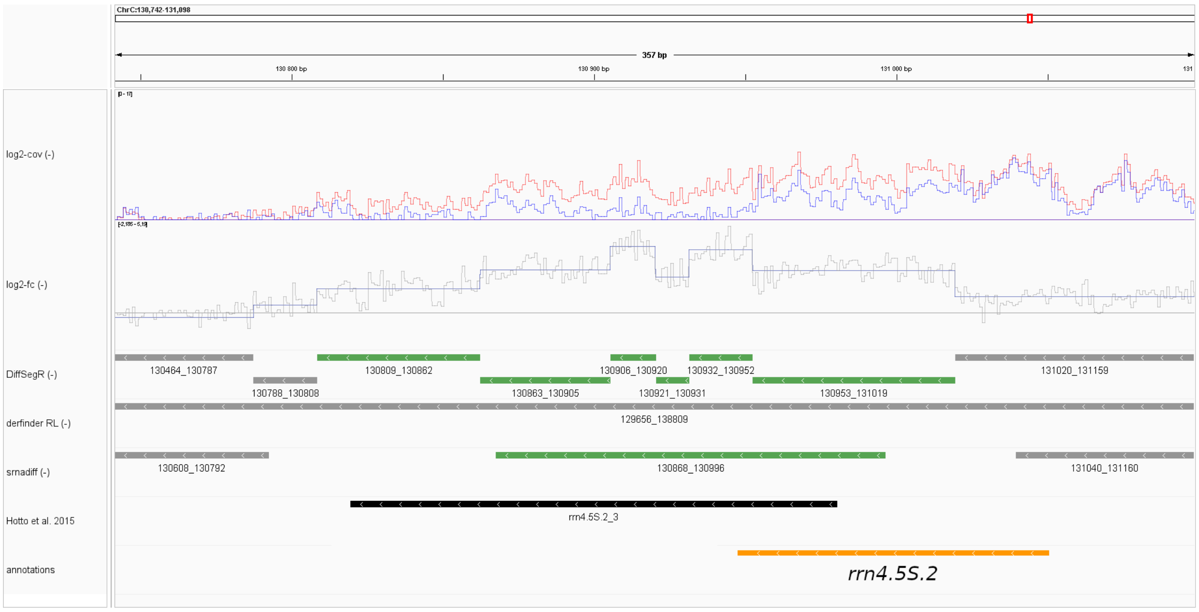


**Figure S26:** Comparison of DiffSegR, derfinder RL, and srnadiff analyses of chloroplast genomic positions 107,549 to 107,905 on the forward strand in the *rnc3/4* dataset. The tracks are similar to those described in Figure S10.

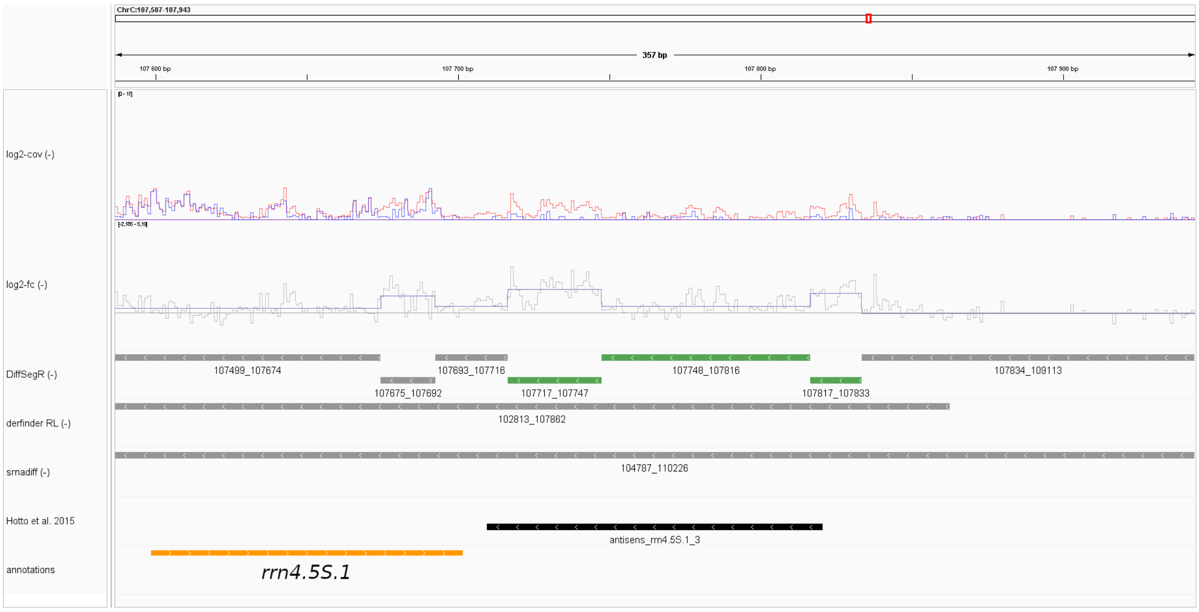


**Figure S27:** Comparison of DiffSegR, derfinder RL, and srnadiff analyses of chloroplast genomic positions 107,587 to 107,943 on the reverse strand in the *rnc3/4* dataset. The tracks are similar to those described in Figure S10.

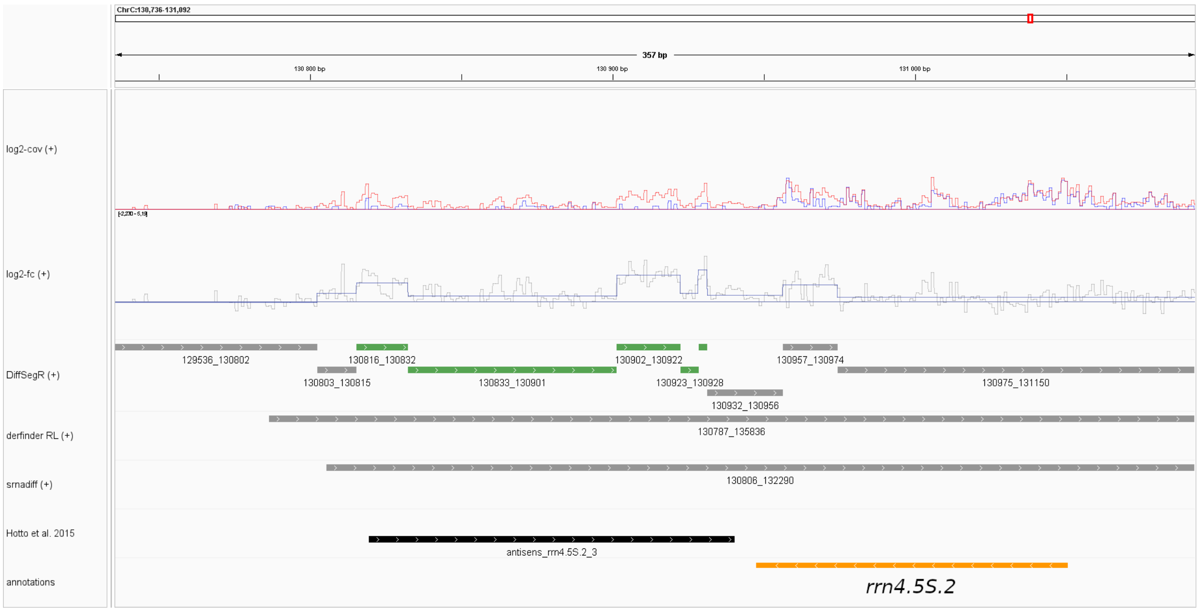


**Figure S28:** Comparison of DiffSegR, derfinder RL, and srnadiff analyses of chloroplast genomic positions 130,736 to 131,092 on the forward strand in the *rnc3/4* dataset. The tracks are similar to those described in Figure S10.

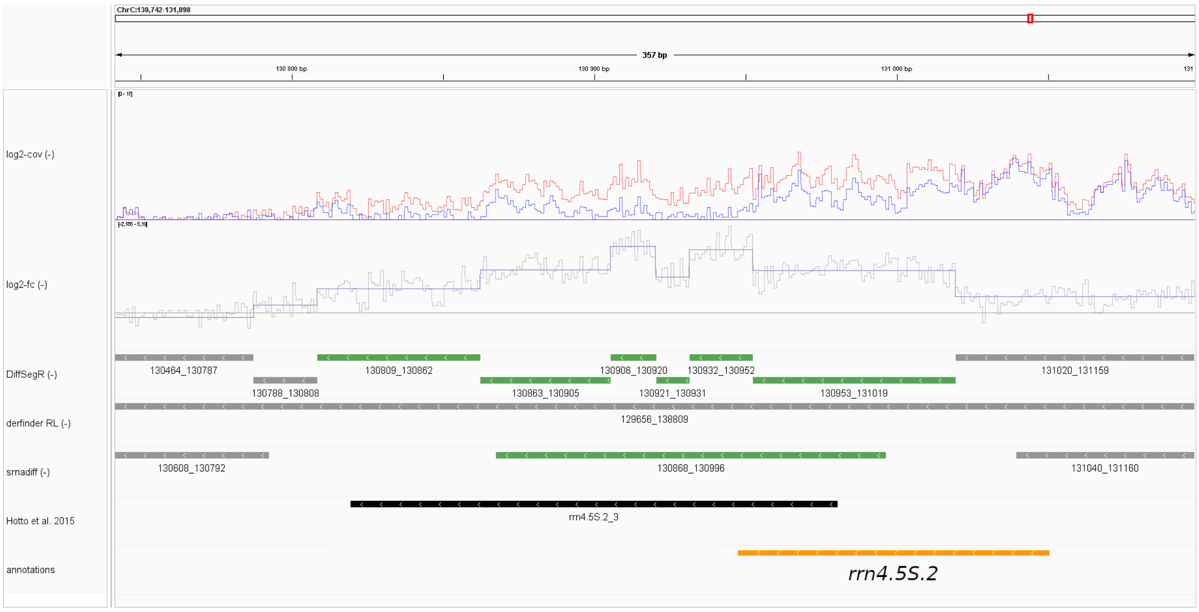


**Figure S29:** Comparison of DiffSegR, derfinder RL, and srnadiff analyses of chloroplast genomic positions 130,742 to 131,098 on the reverse strand in the *rnc3/4* dataset. The tracks are similar to those described in Figure S10.

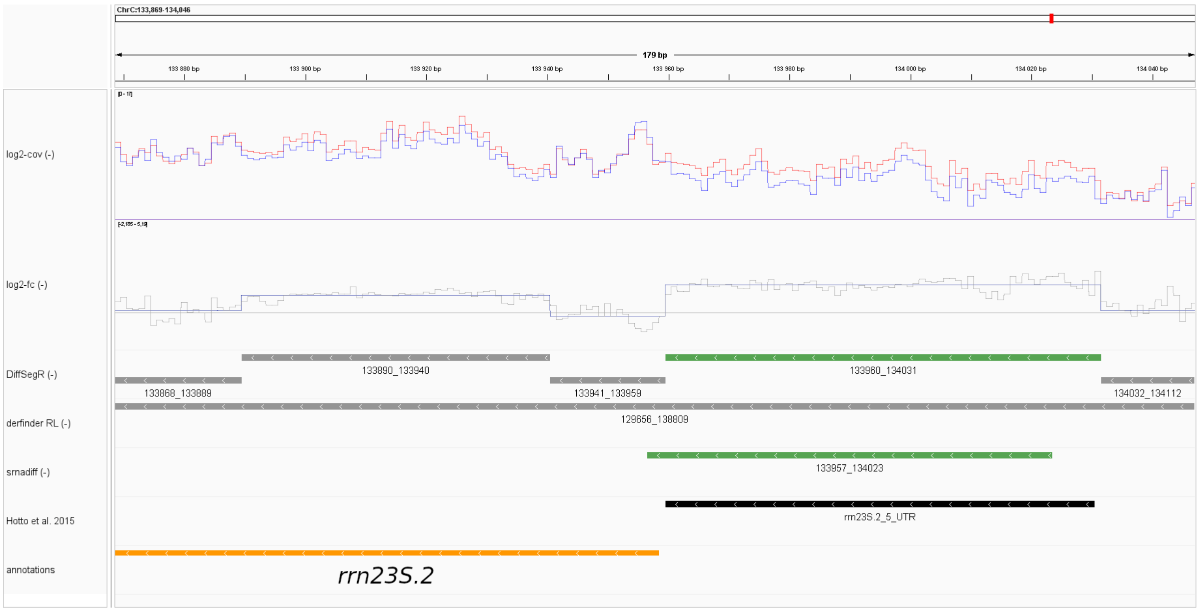


**Figure S30:** Comparison of DiffSegR, derfinder RL, and srnadiff analyses of chloroplast genomic positions 133,869 to 134,046 on the reverse strand in the *rnc3/4* dataset. The tracks are similar to those described in Figure S10.

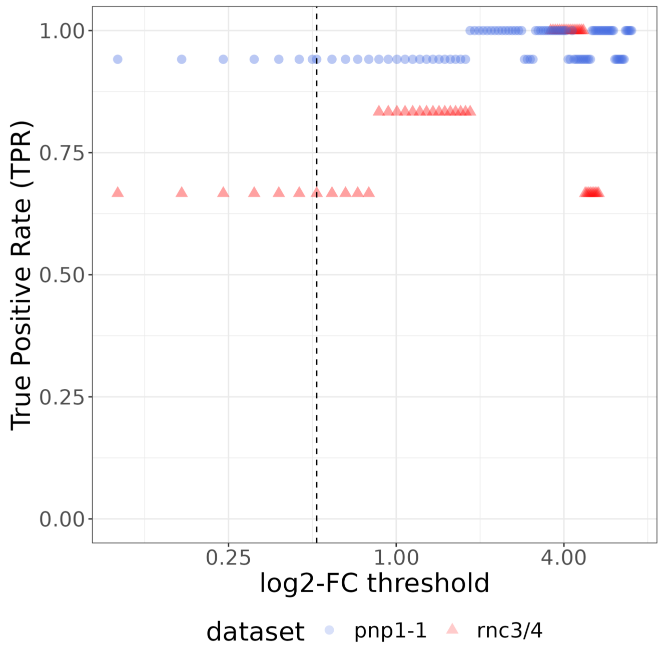


**Figure S31 :** The true positive rate (TPR) of srnadiff on *rnc3/4* and *pnp1-1* labeled datasets as a function of user-defined log2-FC threshold. The black vertical line represents the default log2-FC threshold value (0.5).

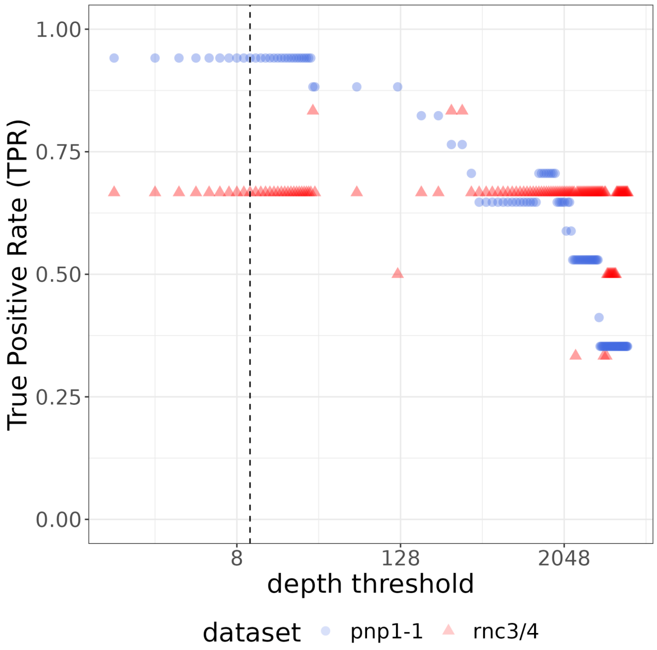


**Figure S32 :** The true positive rate (TPR) of srnadiff on *rnc3/4* and *pnp1-1* labeled datasets as a function of user-defined depth threshold. The black vertical line represents the default depth threshold value (10).

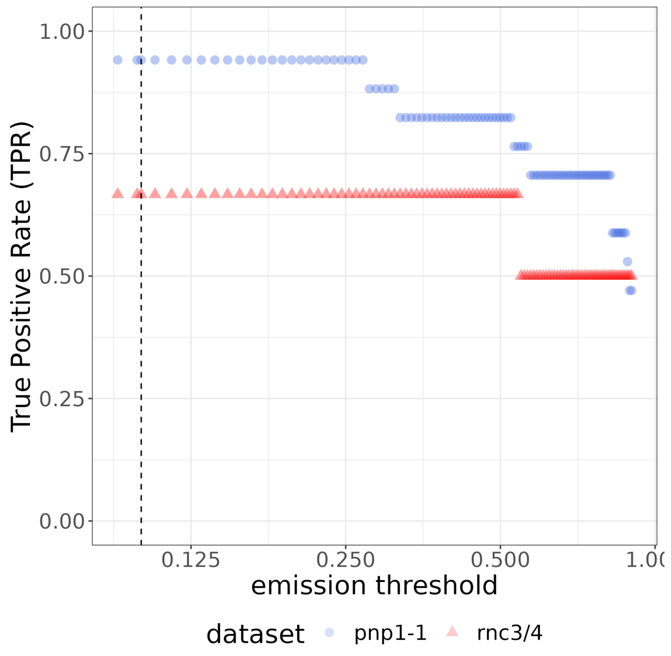


**Figure S33 :** The true positive rate (TPR) of srnadiff on *rnc3/4* and *pnp1-1* labeled datasets as a function of user-defined emission threshold. The black vertical line represents the default emission threshold value (0.1).

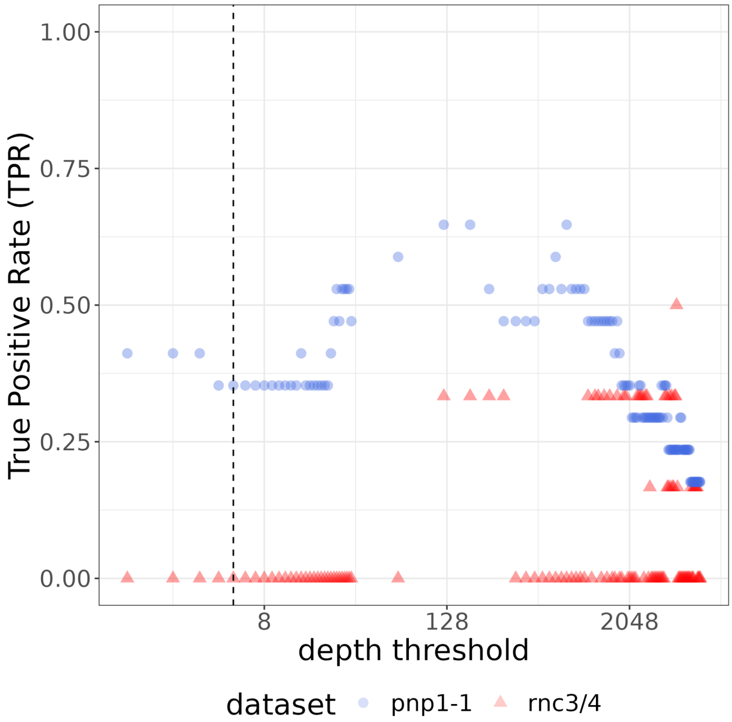


**Figure S34** : The true positive rate (TPR) of derfinder RL on *rnc3/4* and *pnp1-1* labeled datasets as a function of user-defined depth threshold. The black vertical line represents the default depth threshold value (5).

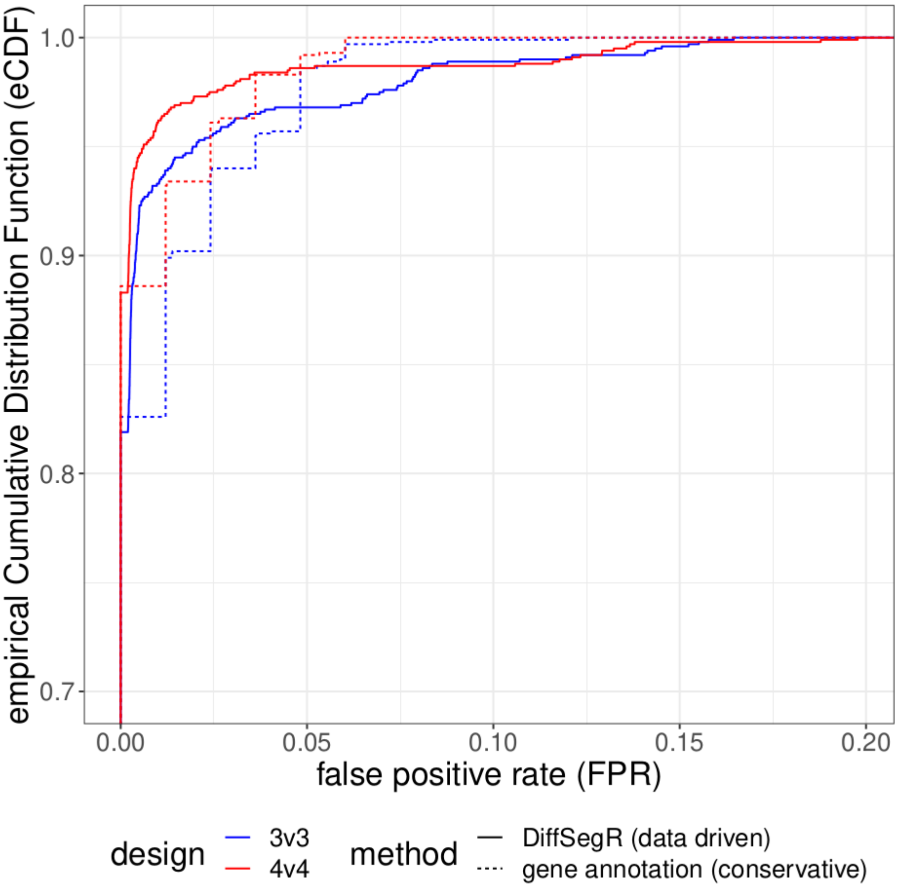


**Figure S35:** Comparison of the empirical cumulative distribution functions (eCDFs) of the False Positive Rate (FPR) from DiffSegR and the Differential Expression analysis within Gene annotations (DGE). The eCDFs of FPRs from DiffSegR (solid curves) and DGE (dashed curves) methods are compared by re-sampling two groups from 10 biological replicates of the same nitrogen deficiency condition in the IDEAs dataset. The figure displays results for group sizes of 3 (blue curves) and 4 (red curves). The eCDF represents the proportion of comparisons (y-axis) with fewer false positives than a specified percentage (x-axis). The eCDF analysis demonstrates that the FPR in DiffSegR results is not inflated compared to the widely-used DGE approach.

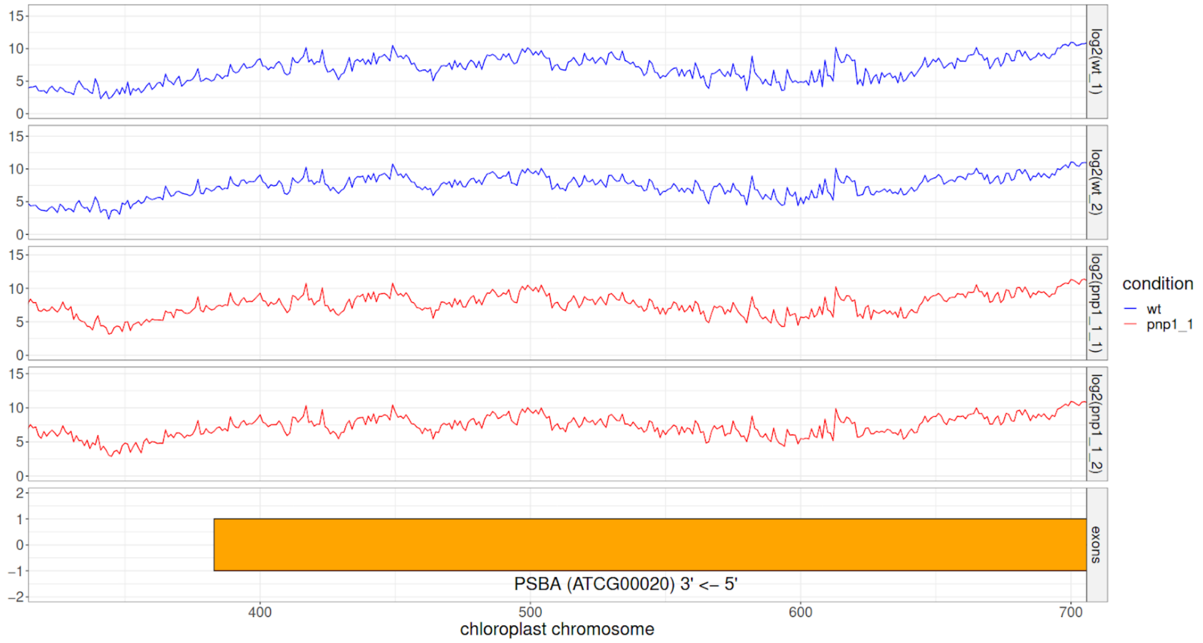


**Figure S36 :** Overview of coverage profiles overlapping *psbA* gene in *pnp1-1* dataset. Coverage profiles exhibit local variations highly reproducible from one sample to another. Local variations could be caused by technical factors that shape the coverage, e.g. 5'/3' bias, PCR bias, GC bias, non-random priming.

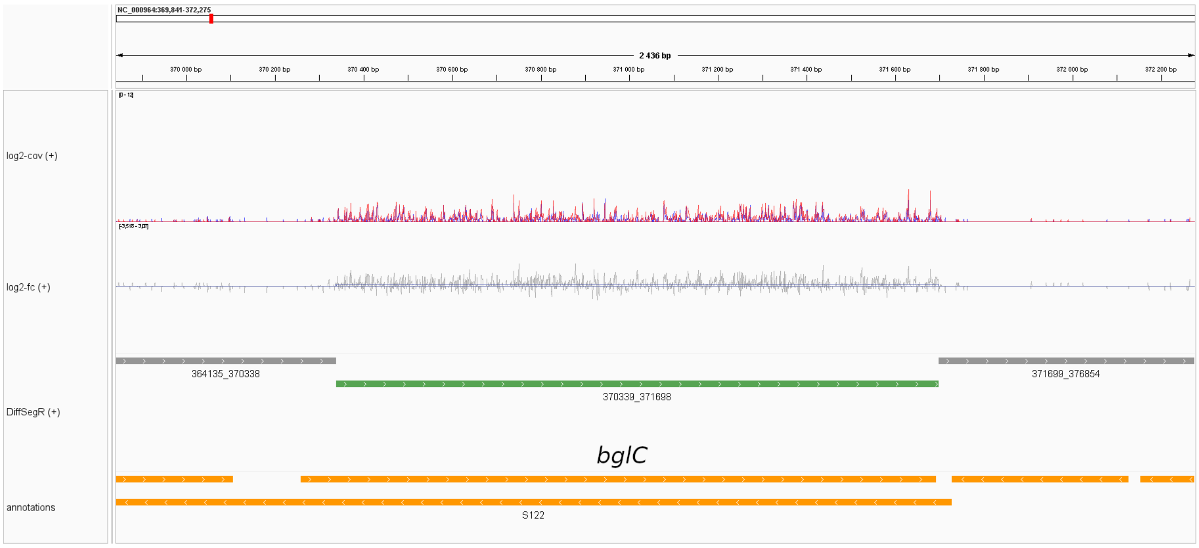


**Figure S37:** DiffSegR analysis of chloroplast genomic positions 369,841 to 372,275 on the forward strand in the $\Delta$*rae1* dataset. The tracks are similar to those described in Figure S10.

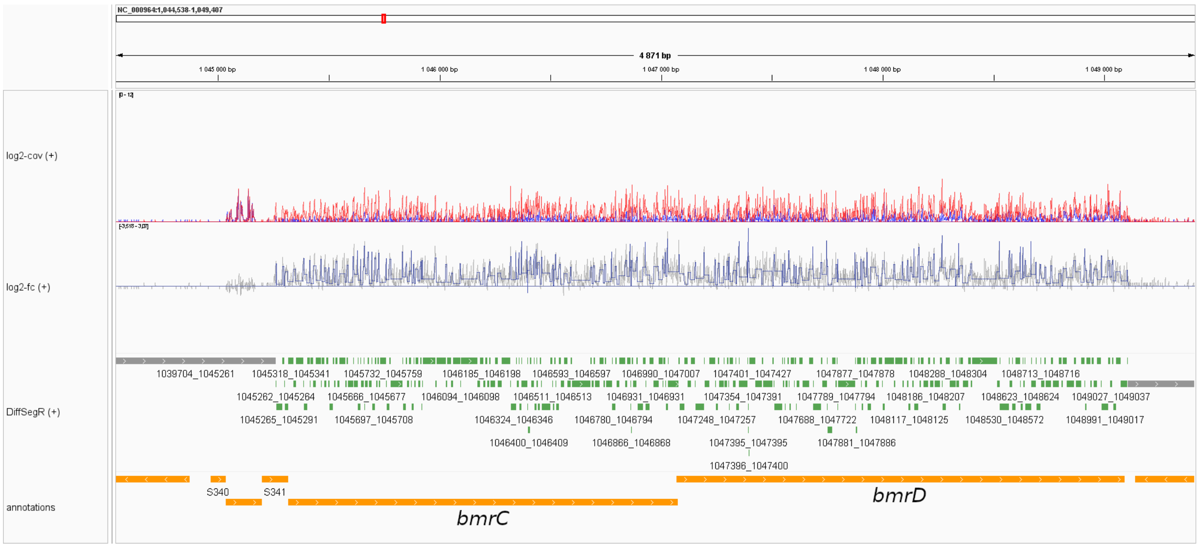


**Figure S38:** DiffSegR analysis of chloroplast genomic positions 1,044,538 to 1,049,407 on the forward strand in the $\Delta$*rae1* dataset. The tracks are similar to those described in Figure S10.

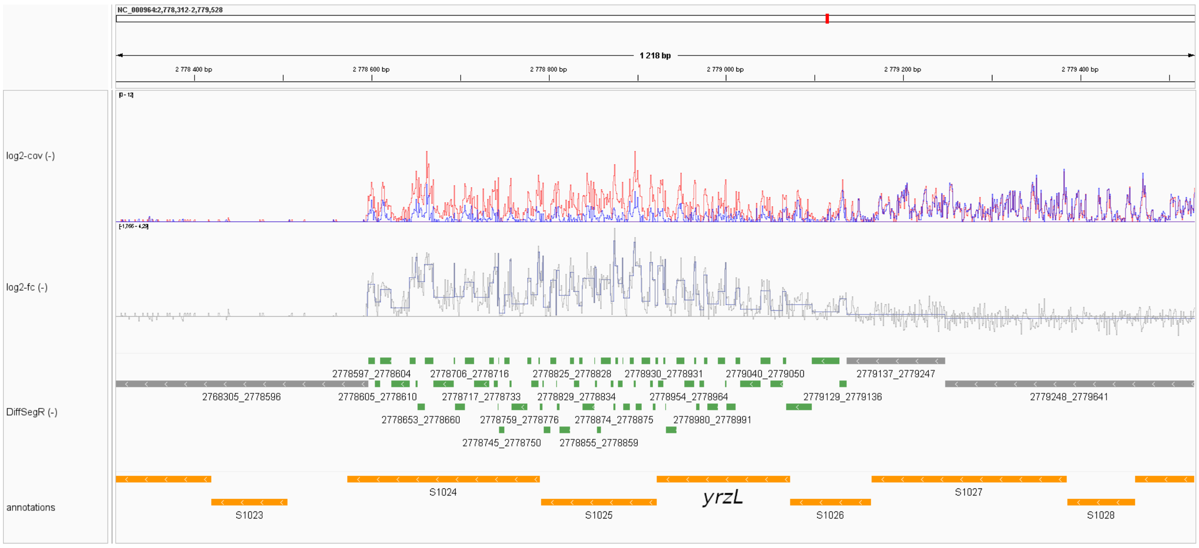


**Figure S39:** DiffSegR analysis of chloroplast genomic positions 2,778,312 to 372,275 on the reverse strand in the $\Delta$*rae1* dataset. The tracks are similar to those described in Figure S10.

### Notes S1 to S5 with Supplementary Table S9 and Supplementary Figures S40 to S44

#### Note S1: Coverage profile Heuristics

We propose four heuristics to compute $Q_{jr}$. The first one takes advantage of the full length reads (Figure S40.A), the second only the 5' ends (Figure S40.B), the third only the 3' ends (Figure S40.C). In previous heuristics, we count the number of elements overlapping each genomic position. For the last heuristic, we compute the geometric mean of the second and third heuristics (as described in the main manuscript).

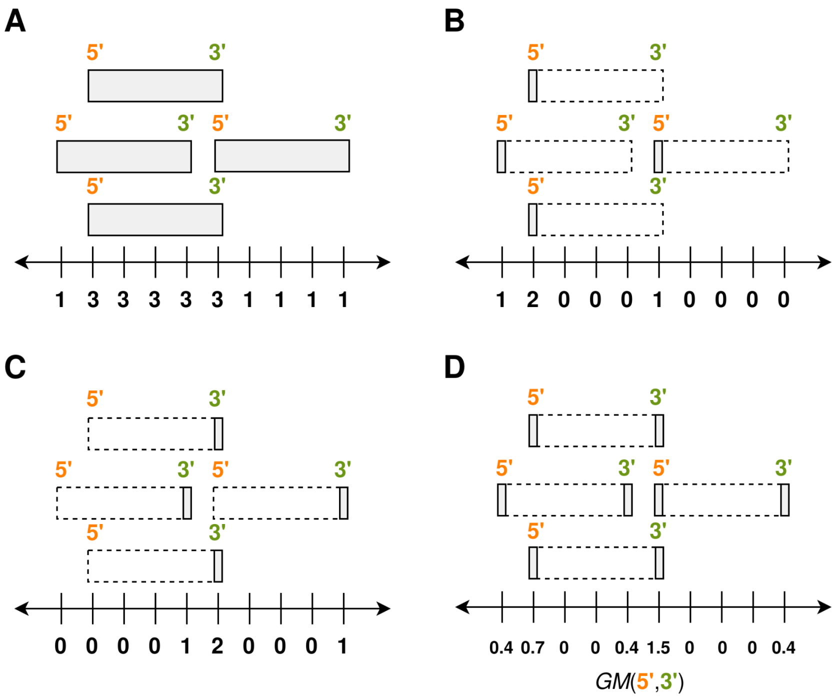


**Figure S40:** Four heuristics used to compute $Q_{j,r}$ from genome mapped reads. In **A** (full length reads), **B** (5' ends) and **C** (3' ends), at each genomic position, elements in gray are summed up to obtain the per-base coverage. In **D** (average) we compute the geometric mean of **B** and **C**.

### Note S2: Full length reads based coverage is more auto-correlated

From a biological perspective, transcription at a particular base is likely to be dependent on transcription at the previous base, but counting a read at all the bases it covers further increases the auto-correlation between counts at neighboring bases. Visually it smooths the profile and gives the false impression that there is little variability. The per-base log2-FC being a transformation of coverages, it is also affected by the auto-correlation. We aimed for a balance between biological consistency and statistical difficulty.

For coverage heuristics, we computed the lag-1 to lag-5 auto-correlation function (ACF) on the per-base log2-FC in both *rnc3/4* and *pnp1-1* datasets. As expected, in *rnc3/4* the lag-5 ACF $\in [0, 1]$ is larger on the per-base lof2-FC calculated on full lengths reads based coverage (0.967) than those calculated on the 5' ends (0.341), the 3' ends (0.379) or the average profile of both (0.483). We observe the same tendency in *pnp1-1* with a lag-5 ACF of respectively 0.989, 0.690, 0.704 and 0.801. Results from lag-1 to lag-4 are available in Table S8. The segmentation model implemented in DiffSegR assumes that counts at every base pair are independent but still have a certain level robust to auto-correlation, certainly not 0.967 or 0.989.

**Table S9:** The per-base log2-FC is affected by the auto-correlation. As empirical confirmation, we computed the lag-1 to lag-5 auto-correlation function (ACF) on the log2-FC per-base in both *rnc3/4* and *pnp1-1* datasets.

| coverage type \ dataset & ACF lag | *pnp1-1* | | | | $\Delta$*rae1* | | | |
|---|---|---|---|---|---|---|---|---|
| | lag-1 | lag-2 | lag-3 | lag-4 | lag-1 | lag-2 | lag-3 | lag-4 |
| full length | 0.998 | 0.996 | 0.994 | 0.992 | 0.994 | 0.988 | 0.981 | 0.975 |
| 5' ends | 0.726 | 0.713 | 0.704 | 0.696 | 0.416 | 0.396 | 0.373 | 0.355 |
| 3' ends | 0.726 | 0.719 | 0.712 | 0.709 | 0.433 | 0.416 | 0.402 | 0.389 |
| geometric mean | 0.820 | 0.812 | 0.807 | 0.804 | 0.532 | 0.518 | 0.504 | 0.491 |

### Note S3: Flanking expressed regions are biased in 3' or 5’ end of reads based coverages

The reads in our libraries have lengths longer than 75 nt. This means that the coverage profiles computed using the 5' end of the reads do not adequately cover the 3' end of the expressed regions, and vice versa for the coverage profiles calculated using the 3' end of the reads. Taking the geometric mean of the two profiles partially resolves this issue (Figures S41-42).

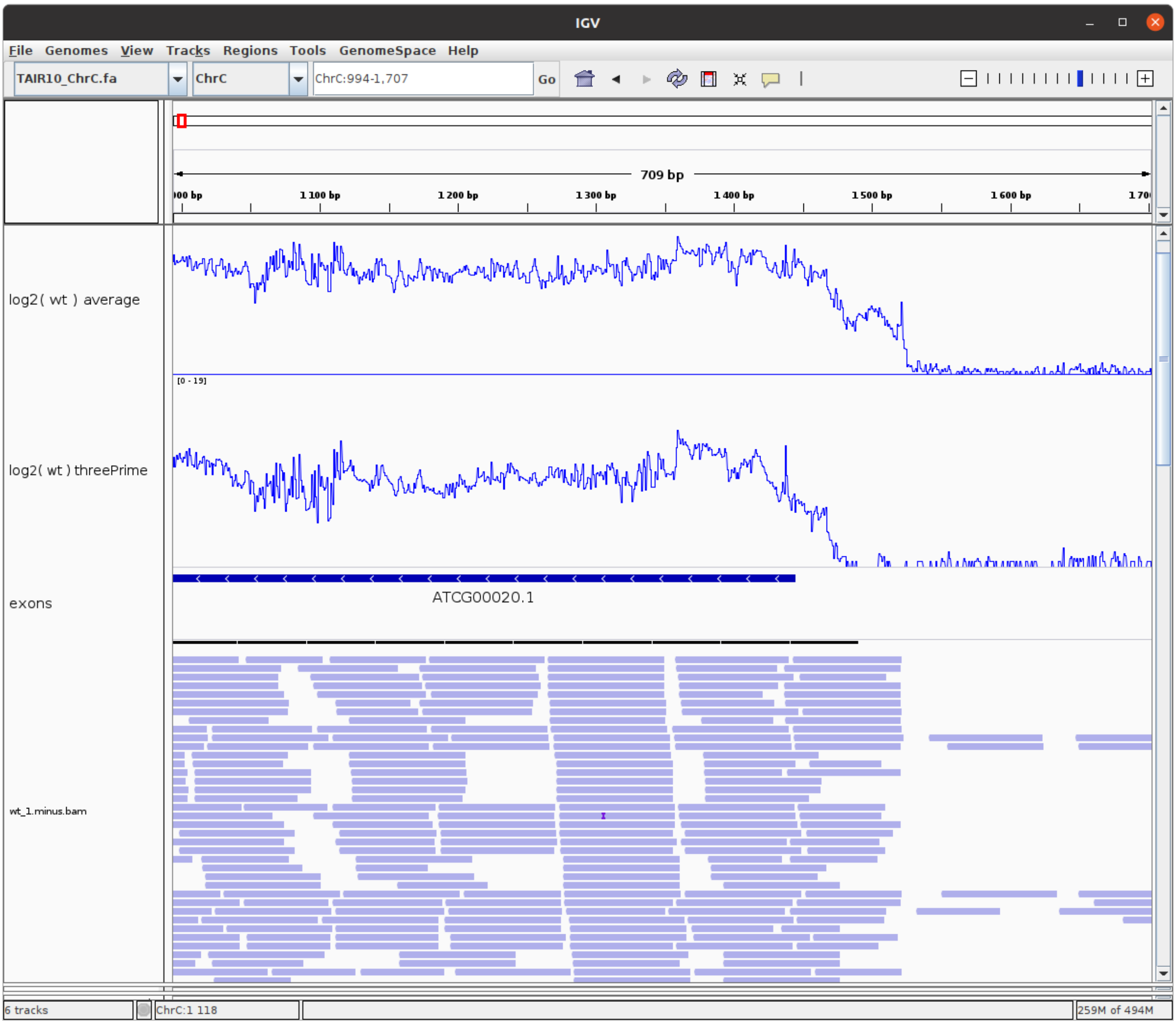


**Figure S41:** 3' ends of the expressed regions are poorly covered in the coverage profiles calculated on 5' ends of reads. Overview of the positions 214 to 927 on the reverse strand of the chloroplast genome. The expressed region corresponds to 3' ends of the *psbA* gene.

The first track stands for the coverage, on logarithmic scale, based on the average of 5' and 3' ends of the *WT* condition in *pnp1-1* dataset The second track stands for the coverage profile calculated on the 5' ends of reads, also on a logarithmic scale, of the same condition. The third track stands for exons boundaries. The last track stands for a sample of mapped reads.

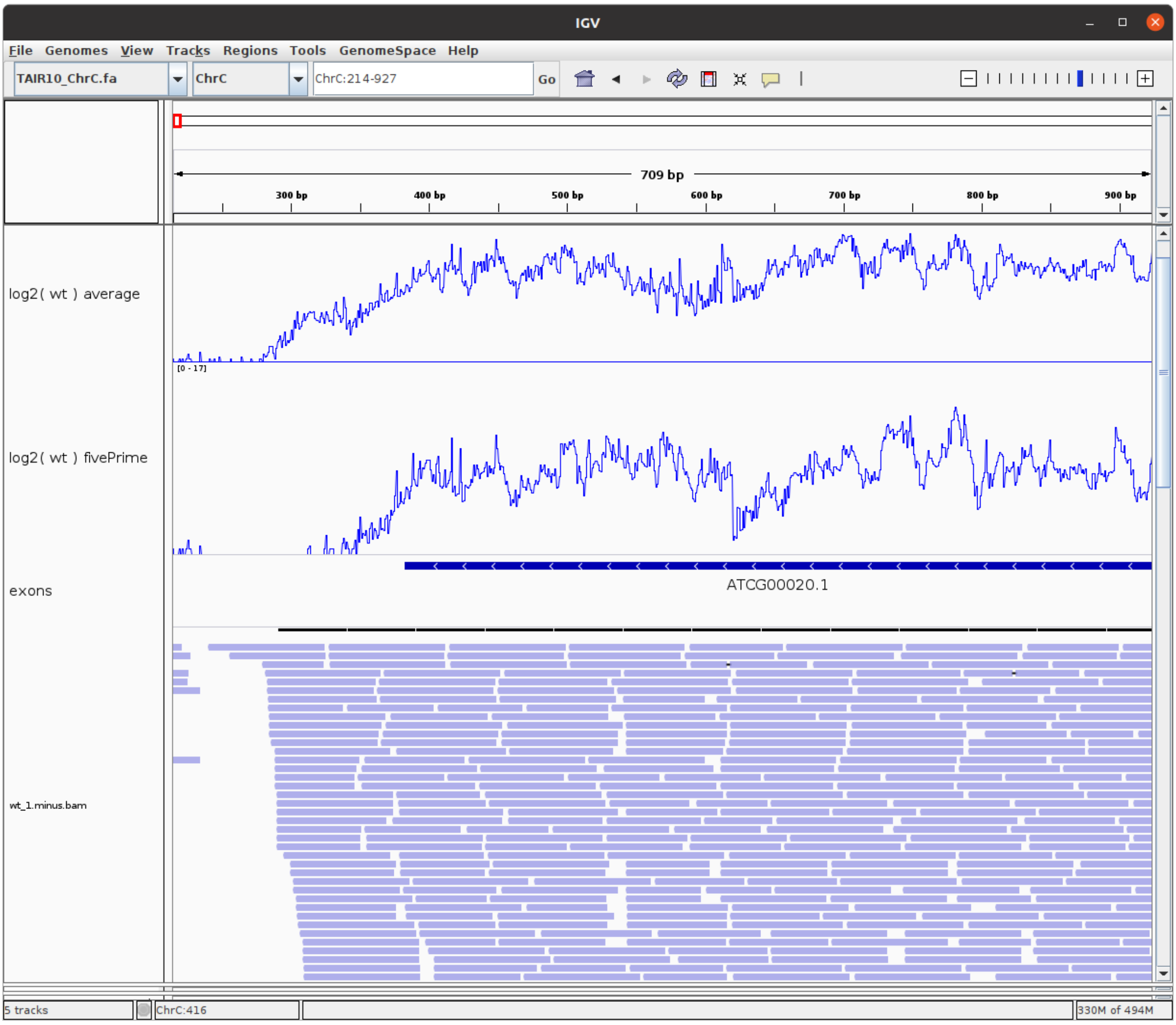


**Figure S42:** 5' ends of the expressed regions are poorly covered in the coverage profiles calculated on 3' ends of reads. Overview of the positions 994 to 1,707 on the reverse strand of the chloroplast genome. The expressed region corresponds to 5' ends of the *psbA* gene. The first track stands for the coverage, on logarithmic scale, based on the average of 5' and 3' ends of the *WT* condition in the *pnp1-1* dataset. The second track stands for the coverage profile calculated on the 5' ends of reads, also on a logarithmic scale, of the same condition. The third track stands for exons boundaries. The last track stands for a sample of mapped reads.

### Note S4: Segmenting in two or three levels merge neighboring differential regions with different log2-FC

In the following paragraph we use a theoretical example to explain the limits of the segmentation models used by state-of-the-art methods to recover differentially expressed regions.

Apart from parseq, which segments the mean of coverages, several other tools (derfinder SB, derfinder RL, srnadiff HMM) use a two-level segmentation with differentially expressed (DE) and non-DE levels, or expressed and not-expressed levels. srnadiff IR uses a three-level segmentation with down-regulated, up-regulated, and non-DE levels. However, we argue that this can be detrimental to biological interpretation. For example, if a gene is up-regulated in condition 2 compared to condition 1 and has an intron retention, a two or three-level segmentation will result in a single large region that effectively combines the up-regulation and intron retention (Figure S43). Additionally, two-level segmentation can merge differential regions with opposite signs of log2-FC, which can also reduce statistical power. A segmentation model that does not make assumptions about the number of levels in the per-base log2-FC should be able to discriminate adjacent differently expressed regulatory events, and this is what we tested in DiffSegR.

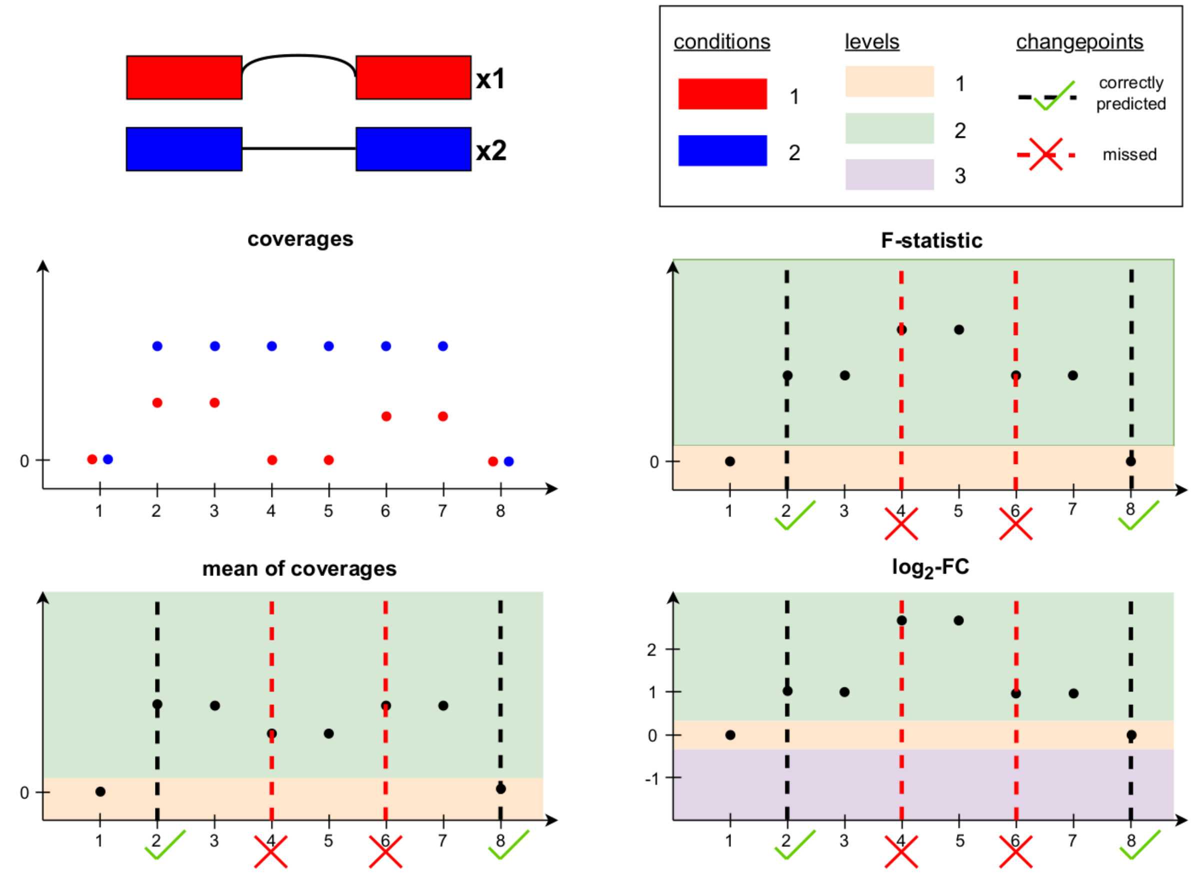


**Figure S43:** Segmenting in two or three levels merge neighboring differential regions with different per-base log2-FC. In this second example the gene is two times more expressed in condition 1 than in condition 2, and in condition 2 the gene also undergoes intron retention. As a result, the log2-FC is higher within the intron than within the exons. Segmenting the mean of coverages in two levels (expressed and not-expressed) merges the up-regulation and intron retention. Segmenting the F-statistic in two levels (differentially expressed (DE) and non-DE) or the per-base log2-FC in three levels (up-regulated, down-regulated, non-DE) also results in the merging of these two events.

### Note S5: Additional changes in the per-base coverage and F-statistic

In the following paragraph we use a theoretical example to explain the limits of the signals used by state-of-the-art methods to recover differentially expressed regions. These limits are assessed on real data in the *DiffSegR better captures the differential landscape* section.

There are various factors that can influence the coverage of a DNA sequence, including the per-base expression, 5'/3' bias, PCR bias, GC bias, and non-random priming. This can result in significant variation in coverage from one base to another. Therefore, it is possible that changes in coverage may not always align with differences in transcription between two biological conditions.

Ignoring any normalization issue, consider bases that follow each other with (non-differential scenario) respectively 1 and 40 counts in both condition 1 and 2 ; (differential scenario) respectively 1 and 40 counts in condition 1 & 3 and 120 counts in condition 2.

There is a noticeable change in coverage between the two bases, yet the log2-FC remains constant and equal to 0 in the non-differential scenario and $\log(3)$ in the differential scenario. These additional changes likely lose statistical power and (in the absence of a post-processing step) make it more difficult to interpret the results biologically, as a single biological regulation will be identified as two or more. Note that, with respect to these two examples, the F-statistic is better behaved yet not perfect. In the first scenario the F-statistics should be equal to its expected value under the null and we should not detect any change between the two positions. However in the differential scenario as power depends on the underlying counts it is likely that we will detect a change between position 1 and 2. The non-differential scenario is illustrated in Figure S44 by the positions 6, 7 and 8. The differential scenario is illustrated in Figure S44 by positions 2 and 3. Given that our primary goal is to identify differences in transcription between two biological conditions, we believe that directly segmenting the per-base log2-FC is a better option, and this is what we tested in DiffSegR.

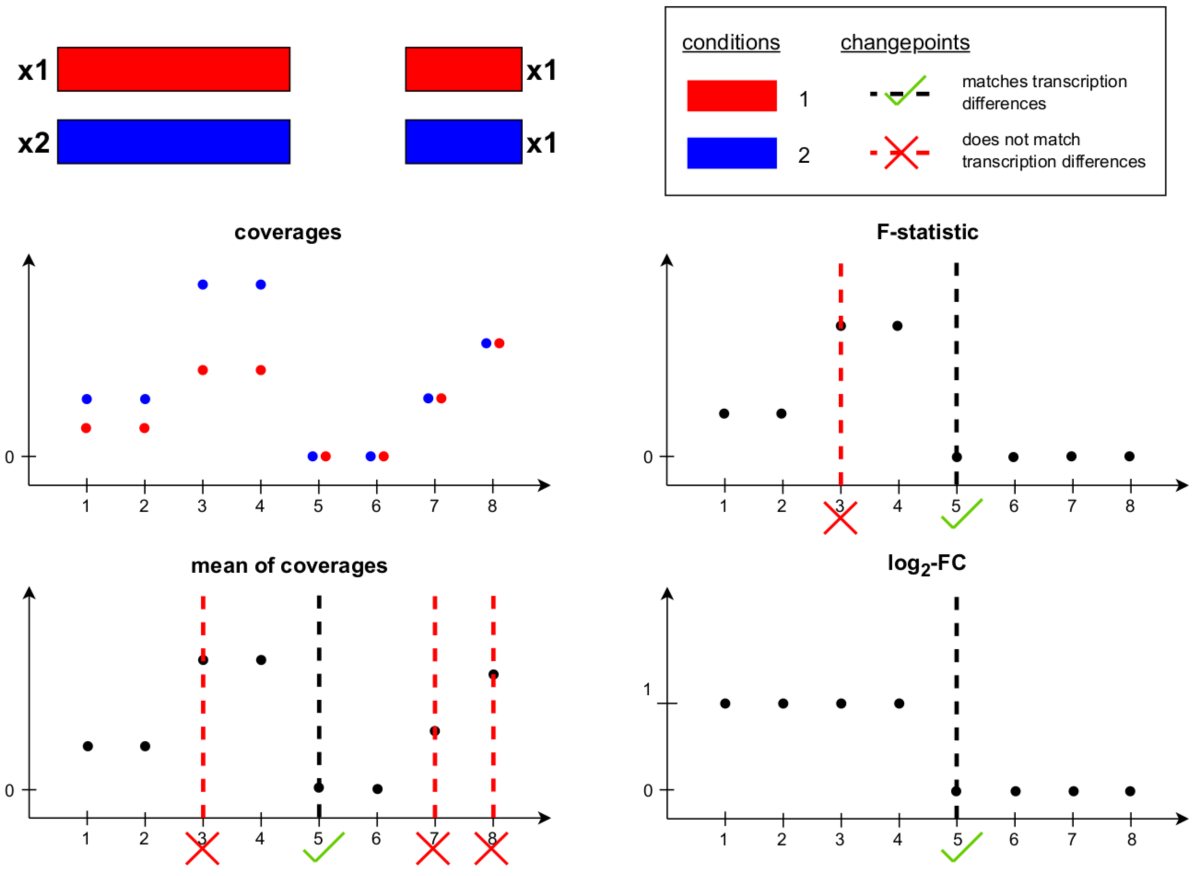


**Figure S44:** Additional changes in the per-base coverage and F-statistic. The first gene is twice as highly expressed in condition 2 compared to condition 1, while the second gene has the same level of expression in both conditions. Segmentation of the mean of coverages, the F-statistic, and the per-base log2-FC results in 4, 2, and 1 changes, respectively. The changes between positions 2 to 3 and 7 to 8 are caused by coverage bias, while the changes between positions 4 to 5 and 6 to 7 mark the end of transcription of the first gene and the start of transcription of the second gene. The change between positions 4 to 5 is the only one that also corresponds to a difference in transcription between the two conditions.
